# Leveling up: improving power in fMRI by moving beyond cluster-level inference

**DOI:** 10.1101/2021.09.23.461354

**Authors:** Stephanie Noble, Amanda F. Mejia, Andrew Zalesky, Dustin Scheinost

## Abstract

Inference in neuroimaging commonly occurs at the level of “clusters” of neighboring voxels or connections, thought to reflect functionally specific brain areas. Yet increasingly large studies reveal effects that are shared throughout the brain, suggesting that reported clusters may only reflect the “tip of the iceberg” of underlying effects. Here, we empirically compare power of traditional levels of inference (edge and cluster) with broader levels of inference (network and whole-brain) by resampling functional connectivity data from the Human Connectome Project (n=40, 80, 120). Only network- and whole brain-level inference attained or surpassed “adequate” power (*β*=80%) to detect an average effect, with almost double the power for network-compared with cluster-level procedures at more typical sample sizes. Likewise, effects tended to be widespread, and more widespread pooling resulted in stronger magnitude effects. Power also substantially increased when controlling FDR rather than FWER. Importantly, there may be similar implications for task-based activation analyses where effects are also increasingly understood to be widespread. However, increased power with broader levels of inference may diminish the specificity to localize effects, especially for non-task contexts. These findings underscore the benefit of shifting the scale of inference to better capture the underlying signal, which may unlock opportunities for discovery in human neuroimaging.

## Introduction

Functional magnetic resonance imaging (fMRI) is one of the most popular techniques available for exploring the living human brain. The bulk of fMRI research is geared towards pinpointing brain areas or networks associated with behavior, traits, and more. When mapping brain areas, researchers typically proceed by estimating a statistic at each of the smallest units of the brain scan, called voxels, then performing inference by pooling information across “clusters” of neighboring voxels. This is more powerful than voxel-level inference when neighboring voxels show sufficient dependence (i.e., share similar properties)—as is the case in fMRI (1)—in part because it avoids multiple testing corrections across thousands of relatively noisy voxels. Cluster-based procedures have also been translated for inference in fMRI brain networks, or functional connectivity; namely, the Network-Based Statistic (NBS; (2)) is used to perform inference on clusters defined as groups of adjacent edges (i.e., “components”).

While performing inference at the areal level accurately reflects some properties of the underlying signal, even decades ago the designers of this approach remarked that more distributed models may better capture the underlying biology (1). Recent work examining larger-than-ever datasets has begun to reveal the widespread dependence in activity across non-neighboring areas of the brain (3, 4). Furthermore, large-scale brain networks that span the brain have been repeatedly identified (5–7). While some have indicated the promise of larger-scale methods for inference (8–10) or prediction (11), a comparison of sensitivity and specificity across levels of inference is lacking. In the meantime, cluster-based inference remains the primary workhorse for typical fMRI studies.

To examine which spatial scale may be most informative for typical studies, we resampled task-based functional connectivity data to empirically compare power across four levels of inference: edges, clusters, large-scale networks, and the whole brain. Critically, designating the full dataset as the population of interest from which subsets are sampled enables us to fully determine “ground truth” effects while preserving the structure of real data. We also designed a “fake” task contrast from resting state data to confirm whether procedures controlled false positive rates as intended. The tools developed here draw upon recent computational frameworks for benchmarking fMRI statistical procedures in large, real datasets (3, 12), including our previous work in the task-based activation context (4).

Altogether, broader levels of inference provided substantially greater power to detect “known” empirical effects. In fact, at our smallest sample size (n=40, almost double the typical sample size of n=25 from (13, 14)), the power to detect an average effect at the edge- and cluster-levels was nearly half that of the network- and whole brain-levels, although this increase in power may come at the expense of reduced localizing specificity (i.e. greater false positives at the scale of connections). Even for the largest sample size measured here (n=120), cluster-level effects still missed effects in more than half the connectome. Only network- and wholebrain level inference approached or exceeded commonly targeted power levels. In the following, we demonstrate how this gain in power is closely related to the widespread nature of “ground truth” effects and other factors, and how controlling the expected proportion of false positives (aka false discovery rate; FDR) instead of the chance of at least one false positive (aka familywise error rate; FWER) can provide substantial benefits in the case of widespread effects.

## Results

We empirically estimated power across levels of inference by resampling functional connectomes derived from the Human Connectome Project (HCP) S1200 dataset (methods summary in **Fig. 1**; see **Methods** for details). Seven procedures were used to perform inference at each level: 1) **edge** via Bonferroni (parametric; (15)), 2) **edge (FDR)** via Storey (parametric; (16)), 3) **cluster size** via NBS; (2)), 4) **cluster TFCE** via threshold-free cluster enhancement (TFCE; (17)), 5) **network** via Constrained NBS (cNBS; (18)) and Bonferroni, 6) **network (FDR)** via cNBS and Simes (19), and 7) **whole brain** via Multivariate cNBS (introduced here). This included a “standard” and a “better” procedure for each level except the whole-brain, for which a single procedure was used; FWER-controlling procedures were used unless “FDR” is indicated. Nonparametric permutation-based procedures were used to control false positive rates except for edge-level inference, which used only parametric procedures (due to feasibility issues with running sufficient permutations for edge-level nonparametric FDR; see **SI Results: Nonparametric edge-level FDR inference implemented in the NBS toolbox**). At each repetition of the resampling procedure, differences between task and resting state connectivity were estimated for a paired sample of subjects. As in (4, 18), resampled data was compared with the full sample “ground truth” dataset and effects detected in the direction dictated by the ground truth effect sign were considered “true positives”. Note that here we define the full dataset as the population of interest, which implies 1) all measured effect sizes are considered exact for the purposes of benchmarking, and 2) effects are expected to be present and nonzero (due to sampling variability) across the full connectome. Generalizability of this paradigm for establishing a ground truth is of course limited, and it might not characterize scenarios in which effects are circumscribed to specific circuits (see **SI Results: Limited accuracy of estimated “ground truth” effects**). Benchmarking experiments were conducted for each of the seven task scans available in the dataset, and three group sizes were chosen for resampling to span from “high typical” to “moderate” sample sizes for this field (n=40, 80, 120; (13, 14)). FWER was also estimated by comparing REST1 and REST2 but shuffling labels for each subject at each resampling repetition.

**Fig. 1.**
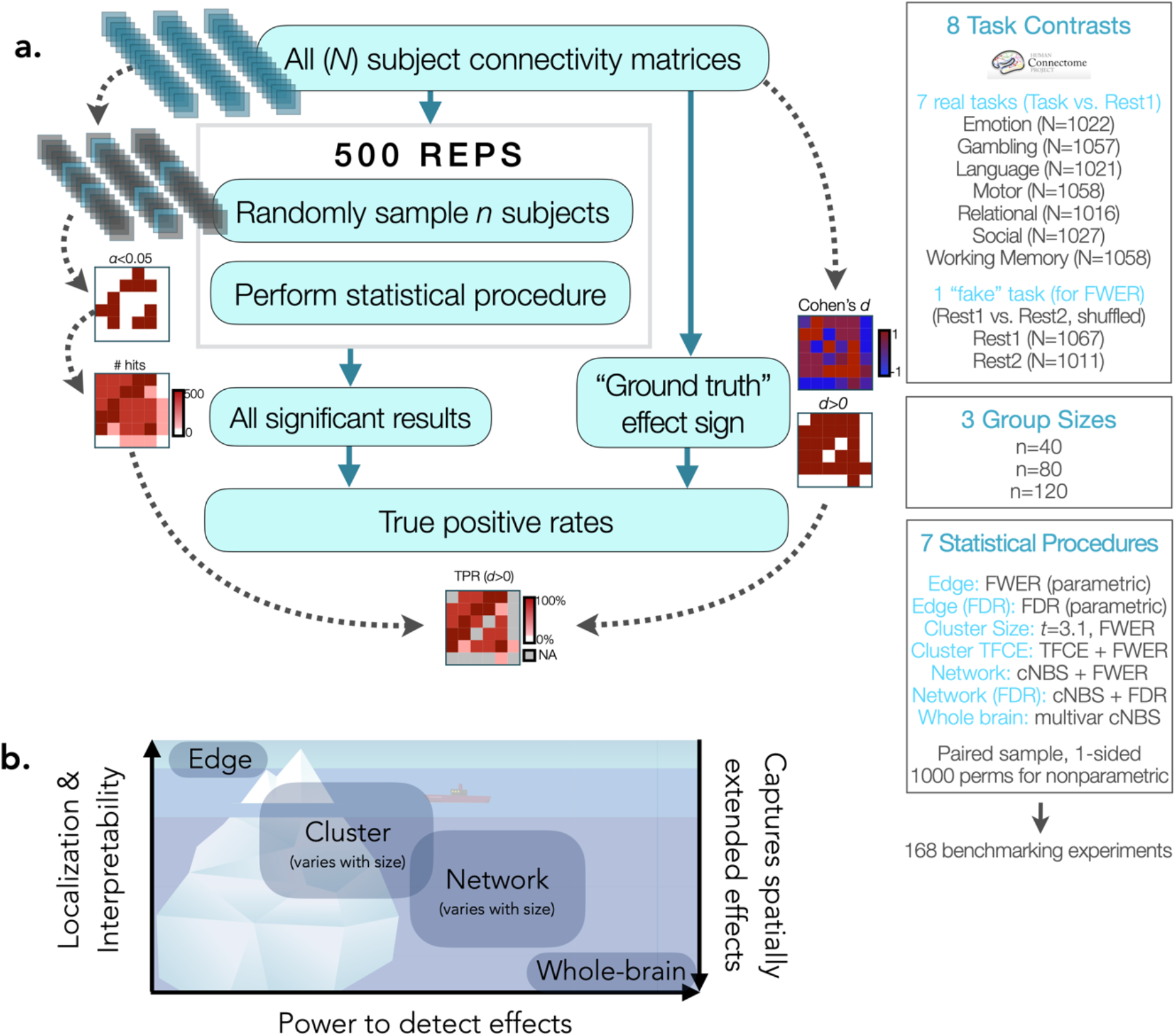
Overview of methods and levels of inference. a) Benchmarking procedure and parameters. b) Tradeoff between ability to localize and interpret results and power to detect effects in typical sample sizes based on the present benchmarking study, along with the extent to which each method captures spatially extended effects.

Overall, the average power to detect a “ground truth” effect was substantially larger for broader levels of inference and FDR control (**Fig. 2a**; **SI Fig. 1a.).** Only network- and whole brain-level approaches attained or surpassed “adequate” power, defined here as the commonly targeted *β*=80% power level. The whole-brain level procedure in particular detected all effects even at the smallest sample size. The other approaches ranged from 10% (edge, n=40) to 50% (cluster TFCE, n=120) average power. The difference between procedures was particularly evident in the smallest group measured (n=40), a sample size yet above average for the field. The gap between approaches decreased with larger samples, although a sample larger than the largest measured here (n=120) would be needed to achieved adequate power to detect the average effect for edge- and cluster-level procedures. However, the choice of error rate under control can change this a bit; the gain in power for FDR controlling procedures relative to FWER controlling procedures was substantial enough that edge-level FDR control resulted in greater power than cluster-level FWER controlling procedures. Finally, it is important to note that there are often sacrifices when increasing power; the increase in power for broader-scale inference and FDR-controlling procedures is expected to come at the expense of spatial specificity and false positive control respectively.

**Fig. 2.**
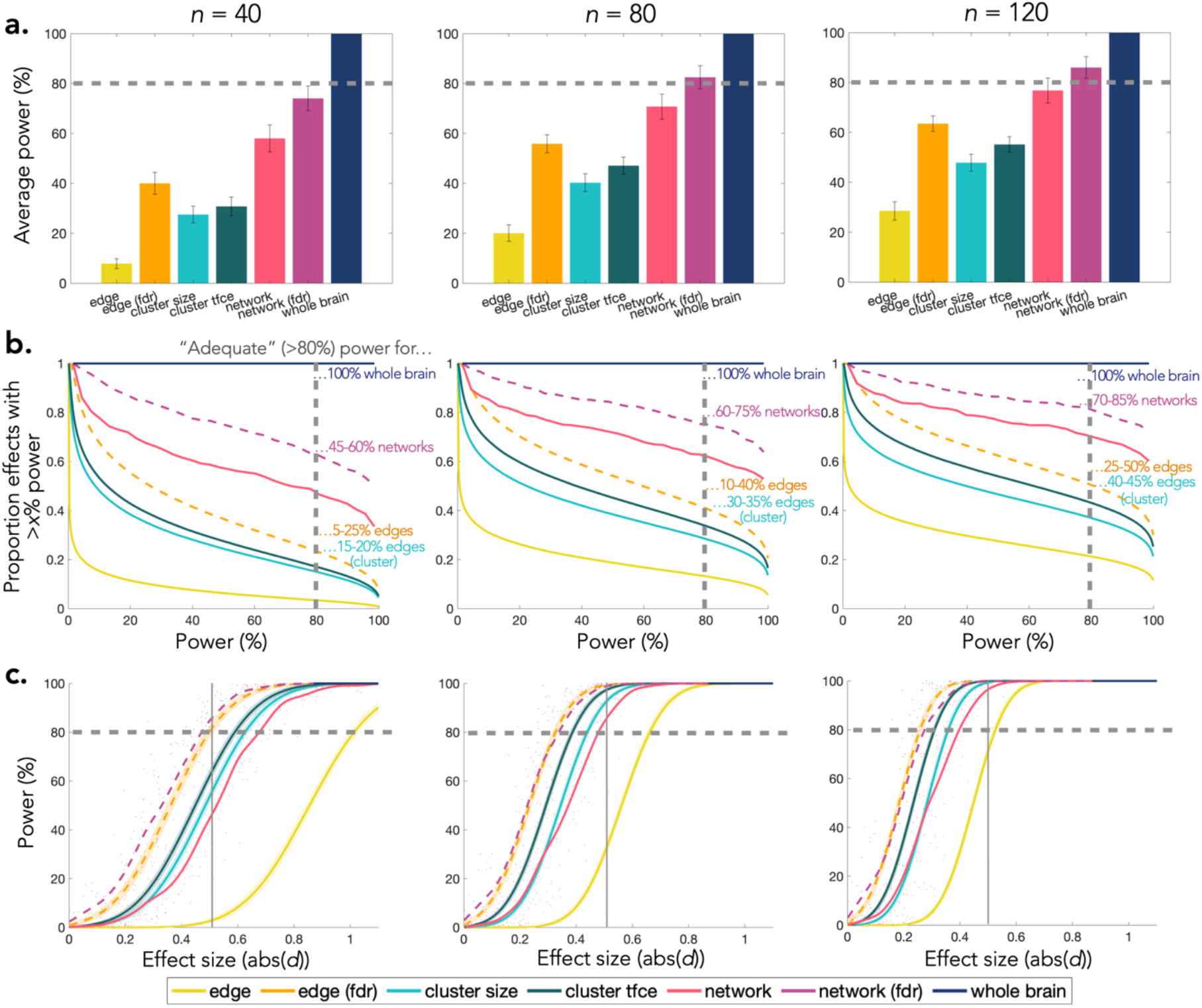
Comparing power across levels of inference. Results are shown for each inferential procedure at three sample sizes, averaged across all seven tasks. a) Average power to detect an effect (e.g., for “edge”, average power across all edges), averaged across all seven tasks for each inferential procedure. Error bars represent standard error of the mean across each of the seven tasks. b) Proportion of effects exceeding each power level across all seven tasks. c) Relationship between power and effect size at each level, with a medium effect size of *d*=0.5 indicated with a thin grey line. Note that distributions of effect sizes are provided in **Fig. 3b**. For all plots, the commonly desired *β*=80% power threshold is indicated by the grey dashed line. Note that power was calculated at the level of each inferential procedure except cluster-level procedures, for which a true positive is defined at the edge level.

We also examined the proportion of effects that were “adequately” powered (**Fig. 2b; SI Fig. 1b**). At the more typical sample size (n=40), adequate power was obtained for a quarter or fewer of edges when using edge- and cluster-level procedures. While this represents a substantial number of edges—a quarter of the connectome is nearly 9,000 edges—it also implies that three-quarters or more of the connectome cannot be detected at desirable rates. In contrast, adequate power was obtained for more than half of the network-level effects at n=40, and, again, all whole-brain effects were consistently detected. Although the gap between approaches decreased with sample size, the more focal approaches remained inadequately powered for over half the connectome with even the largest sample size.

### How the spatial extent of ground truth effects influences power

Several factors contribute to the ability to detect effects. The most important of these is the degree to which the spatial extent of “ground truth” effects matches the inferential procedure used. We note that we estimated nonzero effects for all edges in the connectome, which corresponds with broader scales of inference. To evaluate this decision, we examined the evidence against the null hypothesis across the connectome. On average across tasks, the majority of edges (87%, FDR corrected via Storey; 66%, Bonferroni corrected) and networks (97%, FDR corrected; 82%, Bonferroni corrected) showed significant differences between task and rest with a simple univariate contrast (p<0.05, two-sided t-test; average task effect sizes in **Fig. 3a, SI Fig. 2a,c;** effect size by task in **Fig. 3c**). Furthermore, clusters spanned the whole brain; only a single positive cluster and single negative cluster were found when using cluster-determining thresholds up to a large effect size (i.e., thresholds of |*d*|>0.2, |*d*|>0.5, and |*d*|>0.8 each resulted in only two clusters; **Fig. 3a; SI Fig. 2b**); only using a very large threshold (|*d*|>1.0) produced more than two clusters in a very sparse graph (m=2 positive clusters and m=3 negative clusters). Pooling within networks or across the whole connectome increased effect sizes (**Fig. 3b**, **SI Fig. 2d**), suggesting coordinated activity across widespread brain areas (58% of networks showed medium or larger effect sizes compared with 23% of edges). Importantly, the fact that the majority of effects estimated in the full sample are significant implies that estimated “ground truth” effect signs are meaningful, thus supporting the validity of their use for benchmarking power.

**Fig. 3.**
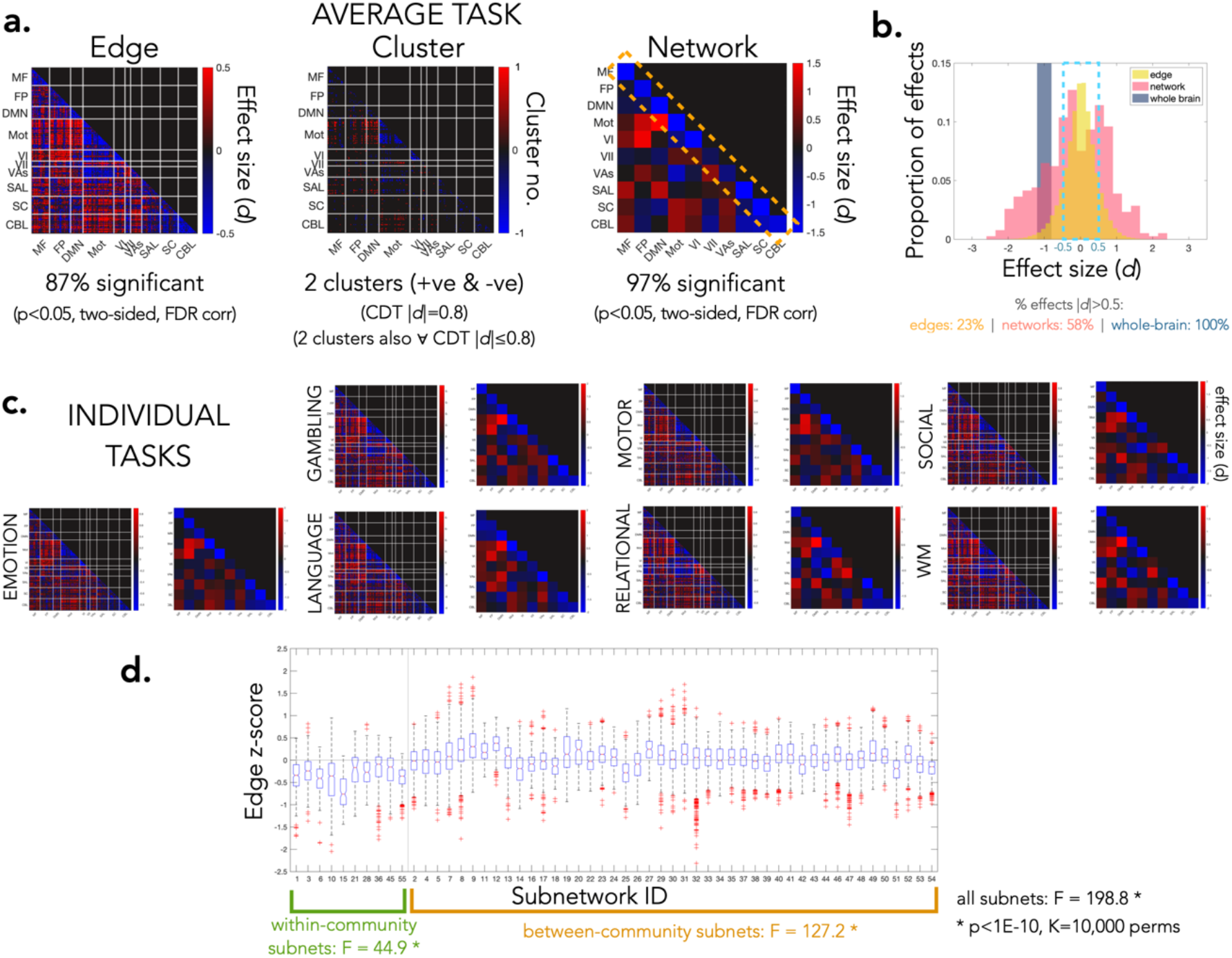
Spatial extent of effects in the full “ground truth” dataset. a) Edge-, cluster-, and network-level effects. The average effect size and average number of significant effects across the seven tasks (task-rest) are shown for the edge- and network-level (*p*<0.05, two-sided t-test, FDR corrected via Storey). For the cluster-level, the average task-rest effect size is used to determine clusters; all edges surviving a cluster-determining threshold of |*d*|=0.8 (i.e., edges with *d*>0.8 and *d*<-0.8) are shown and separate clusters of contiguous edges are counted. Within-community connectivity, generally lower during task than rest, is highlighted by the yellow dotted rectangle. b) Histogram of effect sizes at the edge-level (40 bins), network-level (20 bins), and whole-brain level (i.e., pooled across all edges; 2 bins). c) Edge- and networklevel effects by task. d) Variability between networks. Boxplots show the median (red line), interquartile range (IQR; blue box), and outliers (red whiskers; beyond 1.5*IQR) of edges within each network. F-statistics quantify between-versus within-network variance and significance is estimated by i) shuffling node-community memberships (black), and ii) shuffling edges while keeping within- and between-community structure (green and orange). The vertical line separates within-from between-community networks, and the horizontal line indicates 0.

However, we do not expect simple dependence across the whole connectome; networks also contributed unique information. The original Shen268 networks were significantly more heterogeneous than when randomly shuffling nodes across communities, suggesting that some information may be shared within each network that is not shared across all networks (**Fig. 3d**). Despite notable differences between within- and between-community effects, within-community edges were not interchangeable, and neither were between-community edges; the original within- and between-community networks were more heterogeneous than when shuffling within- or between-community edges respectively (**Fig. 3d**). The Shen268 partition also showed a fair amount of overlap with a partition defined in the HCP with the Louvain method (**SI Fig. 3**). Thus, although pooling within the Shen268 networks is a fairly simple way to account for widespread dependence that is not refined by the structure of the data at hand, it captures some meaningful network-level structure in the independent HCP data (see **SI: Generalizability of the Shen268 partition to the HCP data** for details).

### How the error rate under control (FDR vs. FWER) influences power

As expected, power varied not only with scale of inference but also the error rate under control, favoring FDR (this of course comes at a cost; see **False positives and spatial specificity**). Exploring this further, FDR controlling procedures were more likely to detect an effect of a given size compared with FWER controlling procedures (**Fig. 2c; SI Fig. 1c**). In fact, controlling FDR instead of FWER was like having twice the subjects to detect the same sized effect—for example, for a medium sized network-level effect, power with FDR control at 40 subjects was approximately equal to that of FWER control with 80 subjects (*β*_network_FDR,|*d*|=0.5,*n*=40_=85%; *β*_network,|*d*|=0.5,*n*=80_=84%). Interestingly, FDR controlling procedures offered similar power for the same effect size regardless of level of inference (e.g., for |*d*|=0.5, *n*=40: *β*_edge_FDR,|*d*|=0.5,*n*=40_=80%; *β*_network_FDR,|*d*|=0.5,*n*=40_=85%). For example, edge (FDR) and network (FDR) approaches both had similar power to detect a medium-sized effect at n=40; note, however, that there are much fewer medium-sized effects at the edge level than the network level (**Fig. 3b**). In contrast, there were clear differences between FWER controlling procedures with an intermediate scale of inference actually showing the greatest benefit; cluster-level procedures offered the best power for a given effect size (especially TFCE), whereas the edge-level procedure showed disproportionately low power for the same effect size (e.g., for |*d*|=0.5, *n*=40: *β*_cluster_TFCE,|*d*|=0.5,n=40_=62%; *β*_cluster,|*d*|=0.5,n=40_=53% compared with *β*_edge,|*d*|=0.5,n=40_=3%).

### Spatial bias in effect size and power

The spatial distribution of effects was strikingly consistent across tasks, with decreasing connectivity (towards zero) within-community and between motor and visual communities during task compared with rest (**Fig. 3a,c; SI Fig. 4-5**). These effects are therefore expected to be among the most readily detected during benchmarking. The consistency across task contrasts is primarily because, despite high similarity between task and rest connectomes, rest was distinct from each task in a consistent way. Differences between task and rest were further enhanced by the longer resting scan duration (see **SI Results: Differences between task and rest, and effect of unbalanced scan durations**).

It is already known that edges with larger effects tend to show greater power; we further explored whether some areas showed greater power independent of effect size by examining the residuals of the effect size-power curves (**SI Fig. 6**). This revealed a spatial bias in power independent of effect size—that is, for the same effect size, effects were more likely to be detected in certain areas compared to others. Specifically, edge- (FDR) and cluster-level approaches were slightly more likely to detect effects in subcortical and cerebellar networks at small sample sizes, but this small bias decreased with the larger sample sizes.

### False positives and spatial specificity

It is critical that all inferential procedures achieve expected control of false positives. All procedures, including those designed for FDR control, are expected to control FWER in the weak sense (i.e., when the null is true everywhere; (20). Indeed, all procedures attained FWER below the upper bound of the expected 95% confidence interval (estimated FWER 95% CI = 3–7% for 500 repetitions; (21, 22)) during the “fake” task designed to generate null effects (**Fig. 4**). While some procedures that were more powerful also had higher FWER (notably networklevel procedures), the gain in power for broader scale inference was not solely due to more permissiveness—no false positives were observed for the whole-brain inferential approach despite its 100% power. Also differing from the power results, there did not appear to be a consistent change in FWER with sample size across approaches, and no bias in the spatial distribution of false positives was observed (**SI Fig. 7**). There may be some room for improvement in the edge- and cluster-level approaches, which appeared overconservative (defined as nearing or falling below the lower bound of the 95% confidence interval for FWER). The relationship between sample size and FWER was unclear, but cluster-level approaches may be most overconservative at low sample sizes.

**Fig. 4.**
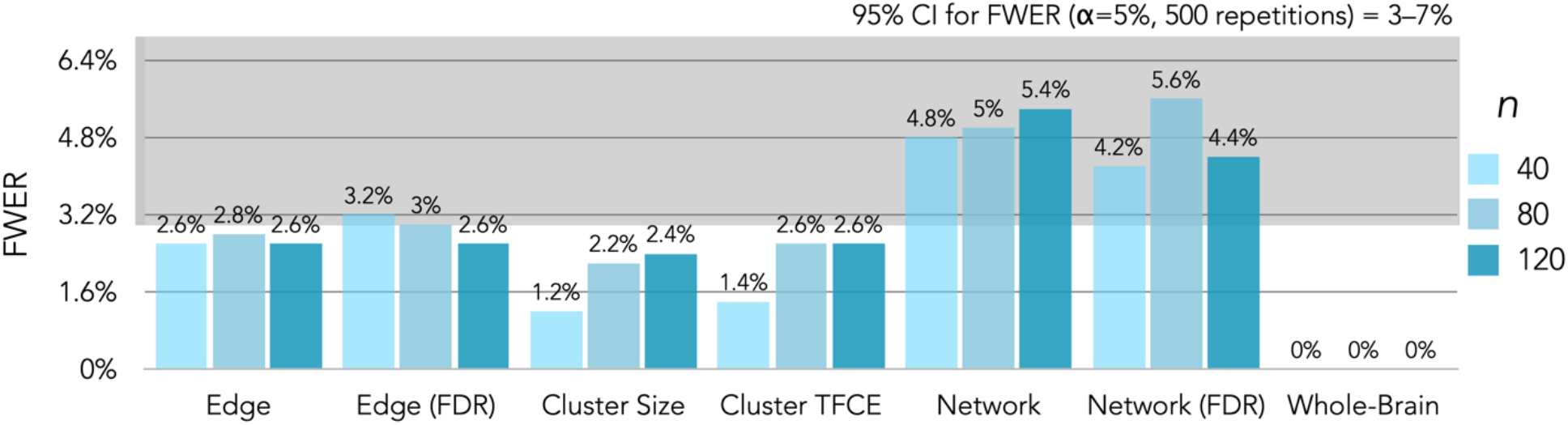
All procedures achieve valid FWER control. The expected 95% confidence interval for the FWER is highlighted in grey. Valid control is defined as limiting FWER below the upper bound of this interval. All procedures beside network-level procedures were over-conservative, defined as nearing or falling below the lower bound of this interval.

This evaluation provides an important but preliminary understanding of the specificity of these procedures. This only included an assessment of weak FWER control (i.e., true null everywhere), but it is important to note that, by definition, FDR controlling procedures do not attain strong FWER control. The implication is that when there are true effects in the network, FDR controlling procedures may permit additional false positives compared to strong FWER controlling procedures in order to achieve the increased power above. It is also important to understand the extent to which procedures provide spatially precise inferences (i.e., localizing power). In general, broader levels of inference decrease spatial specificity unless the spatial extent of the inference matches the spatial extent of the underlying effect. While spatial specificity is not directly assessed here, the ground truth effect size maps suggest that effects are indeed widespread but not perfectly matched to the network-level inference; thus, we expect network-level approaches to provide a fair level of spatial sponeOecificity. **Fig. 1** shows the expected tradeoff between spatial specificity and power. Although here we find that task-based effects tend to be widespread, we remark that alternative ground truth frameworks could be considered in which effects are circumscribed to relatively small proportions of the network. In such cases, edge or cluster level inference may be desirable, particularly if the goal is to provide high localizing power.

## Discussion

Despite the popularity of focal inferential procedures, the present work empirically demonstrates that simple procedures that account for broad-scale dependence better reflect the spatial extent of effects and thus substantially improve power. Together with recent work in large task-based activation datasets (3, 4), these results suggest that cluster-based inference may only reveal the tip of the iceberg of true widespread effects in typically sized task-based studies. These results suggest that leveling up may be an optimal path forward for many typical studies that need to prioritize detection power over localization power.

### Implications and recommendations for functional connectivity inference

Shifting to a broader level scope stands in contrast with a conventional focus on localization first. Indeed, some have suggested that users employ more stringent cluster-determining thresholds specifically to obtain more focal inferences (23). However, the present results challenge the utility and meaningfulness of standard focal inference. First, effects were found to be widespread for a variety of tasks. It seems unlikely that an organ as complex as the brain would have areas wholly uninvolved in many studied cognitive processes. Second, typically sized studies are not well equipped to detect any focal effects that may exist. On the other hand, combining information across the whole brain can also hinder interpretation and utility, e.g., in determining targets for intervention. There is certainly a tradeoff between power and spatial specificity, though it remains unclear which spatial extent is most useful or biologically meaningful (in the sense of reflecting the underlying signal for inference). We believe that a network-level inference approach like cNBS, which leverages widespread dependence while retaining unique network-specific information, is a step in the right direction.

Other approaches that employ pooling and/or leverage multivariate information will offer similar advantages. This naturally points towards multivariate statistics and machine learning, which are both growing increasingly common in fMRI (24). Deciding when to treat variables as independent features or pool them requires some theoretical justification. Simple pooling, as used for cNBS, assumes that pooled variables are random realizations of a shared underlying effect (i.e., “redundant” to an extent). This is useful if noise is expected to be relatively independent across variables within the pool and thus “averaged out” to better estimate the shared signal. This is a fairly simplistic picture of dependence, and it may be more useful to draw upon the multivariate pattern of independent features across the brain. Indeed, we showed significant variability between subnetworks, suggesting they offer independent information. We also treated networks as contributing potentially independent information in multivariate cNBS. One may also explicitly model dependence and independence between variables, e.g. through factor analysis. Either way, smaller studies tend to miss existing effects and/or underestimate the spatial extent of effects, and would benefit not only from combining widespread information but also from leveraging a priori information to guide how effects are combined. This does not preclude finer-grained analysis; in fact, the designers of cluster-level inference recommended that studies with an unknown profile of effects start with set-level inference and illustrated that a step-down approach to identifying increasingly localized effects should not increase the FWER (1). This has recently been formalized and translated for neuroimaging via the “All Resolutions Inference” framework (25). In addition, once existence of an effect is broadly established, more data can be collected for the purpose of localization.

Critically, FDR-controlling procedures had substantially greater power than FWER-controlling procedures, such that more focal inference when used with FDR actually outperformed broader-scale inference when used with FWER. FDR control is beneficial when one is willing to admit more false positives to obtain more true positives, and should control FWER when the null is true everywhere (20). The resulting decrease in spatial specificity may be a reasonable sacrifice in the context of distributed fMRI effects in the present study. The field has highlighted the potential power gains for FDR over FWER (26) and recently, when the field reckoned with invalid FWER control for popular parametric cluster-level inferential procedures, a follow-up analysis suggested that results would still hold if studies had controlled FDR (27). Whether the lagging popularity of FDR is due to convention, accessibility of tools, or other reasons, the present results underscore an opportunity for more FDR-based tools.

### Implications for data beyond functional connectivity

This study is motivated in part by observations from task-based activations; therefore, it is only fitting that we expect implications for that context as well. While many typically sized studies demonstrate results in localized “blobs”, larger studies show widespread activations across the brain (3, 4). Thus, larger scale inference, perhaps also using resting state communities that capture dependent activity, is expected to also benefit activation-based studies as well.

Outside of neuroimaging, much of biomedical research also struggles with maintaining power given high dimensional, dependent data with small univariate effect sizes. Many other modalities in neuroscience simultaneously record many signals (e.g., EEG) and thus also reckon with spatially dependent observations. Genetic methodologists are also exploring approaches that capture widespread effects (28, 29) and translating methods to neuroimaging (11). Since we all face similar issues, it may be fruitful for neuroimaging to examine methods for evaluating and capturing dependence in other areas, and perhaps methods developed in neuroimaging may in turn have relevance to those areas.

### The nature of the signal in the connectome

How the brain functionally reorganizes during task and rest is a major topic of study. As has been shown before (30), we observed that rest and task connectivity were highly similar. However, whereas paired differences between tasks were relatively unique, rest was distinct from each task in a consistent way; task connectivity was generally lower within communities and between motor and visual communities. Previous reports have also showed that integration is greater during task than rest (31, 32) and increases with cognitive demand (33), suggesting that more cognitively complex tasks require more collaboration across systems performing unique functions. These results together illustrate the divide between cognitively demanding task states and less demanding rest, which adds nuance to the common notion that rest explores the full repertoire of task-relevant states. Since contrasting task with rest may reflect general demand-specific rather than cognition-specific effects, investigations may benefit from a more task-relevant reference—perhaps a task with similar cognitive demands, or an estimate of intrinsic connectivity that includes task (e.g., (34). Importantly, task-rest differences were enhanced by the longer resting scan duration and care should be taken to account for differences in the scan duration that can bias contrasts towards the longer (typically rest) scan. Overall, the distinctiveness of rest, taken alongside findings that rest may contain less phenotypically relevant information (35), points towards the need for increased scrutiny when using rest in connectome-based analysis.

More generally, the spatial distribution and magnitude of effects are expected to vary substantially with study design, statistical models, covariates, and more. Brain-behavior associations in particular appear to be an order of magnitude smaller than the present task-based effects; perhaps due to weaker or more heterogeneous effects, differences between edge- and network-level results were not apparent for such associations (36). Thus while we expect results to generalize to reasonably similar study designs, the spatial extent of effects and optimal level of inference for study designs that differ substantially is an open question.

### Open questions for fMRI inference

Pooling within predefined networks is a relatively simple procedure for harnessing the known dependence between widespread areas, and the present study is intended to illustrate the extent to which this improves power. There are a number of ways cNBS and multivariate cNBS can be adapted to more accurately reflect expected effects, from how the partition is estimated *(a priori* from a particular dataset, from the data at hand, etc.), to how dependent data is aggregated (average univariate statistics, average timeseries, across multiple spatial scales, etc.), and more. A major consideration is the yet-to-be determined balance between leveraging larger datasets to help inform smaller studies while also respecting the unique properties of the study at hand, since networks reconfigure with a number of study-specific characteristics (e.g., (37)). Typically sized studies are expected to benefit the most from an *a priori* partition since network characteristics may be more robustly estimated from larger datasets. Making things difficult, there are numerous network definitions currently available and which to use for this partition is an open question. Fortunately, the core components of several major “resting state networks” are moderately reproducible (7) and thus we anticipate that many commonly used network definitions or the universal taxonomy described in (7) may provide a reasonable starting point. One can also gain power by constraining analyses in other ways. It may be advantageous to take a more hypothesis-driven approach to limit the networks under evaluation, preferentially omitting select between-community networks since more tests are conducted between-than within-community.

Finally, as alluded to above, evidence increasingly points towards the trivial nature of a mass null hypothesis in the context of widespread, multivariate effects. A useful alternative may be a Bayesian framework for characterizing the strength of effects and establishing large data-informed priors. Our larger datasets may offer some insights for building these priors; more about the nature, structure, and limitations of the effects estimated here is discussed in the **SI Results**. Overall, the present results are just a demonstration and a starting point, and merely scratch the surface of what possible inferential procedures can be used to account for dependence and combine information across the brain.

## Conclusion

Cluster-level inference enabled many important discoveries when the field was first exploring tailored procedures for more powerful inference beyond the voxel level. Today, larger-than-ever datasets and computational resources have facilitated the evolution of statistical procedures based on a more complete understanding of their accuracy and the nature of the signal. Here we highlighted one important avenue towards continued improvement: the use of procedures designed to capture the widespread spatial effects across the brain, which provide a desperately needed boost in power for typically sized studies. More generally, the impressive rise in openly available datasets in the field presents an opportunity to better characterize elements of the signal to inform inference for more typical studies. As we learn more about data we grow better equipped to build tools tailored to the underlying signal, which, in turn, leads us closer to more robust and reproducible findings in neuroimaging.

## Methods

Functional connectivity data derived from the Human Connectome Project (HCP) S1200 release were used to estimate power and FWER. Minimally preprocessed data provided by the HCP were further processed to regress noise and obtain “connectivity matrices” of the z-scored Pearson correlations between the 268 nodes of the Shen268 atlas (38).

168 “experiments” were used for benchmarking seven inferential procedures (below) at eight task contrasts and three group sizes (n=40, 80, 120). At each experiment, paired sample, onesided tests were used to contrast 7 “real” tasks versus rest (tasks = emotion, gambling, language, motor, relational, social, working memory) or one “fake” task versus rest (contrast between two resting runs, shuffled for each subject). 500 repetitions were used for each nonparametric procedure. For power calculations, discoveries from resampling the real tasks were compared with the full sample “ground truth” dataset (N=1021–1058, depending on the contrast) to determine whether effects were detected in the direction dictated by the ground truth, analogous to one-sided tests conducted in the “correct” direction. For FWER calculations, all resampling discoveries from the “fake” task were classified as false positives and the rate of at least one false positive per repetition was measured.

The seven procedures used to perform inference at each of the four levels were: 1) **“edge”**: a parametric procedure for FWER correction (Bonferroni procedure), 2) **“edge (FDR)”**: a parametric procedure for false discovery rate (FDR) correction (Storey procedure), 3) **“cluster”**: the NBS a method for cluster-level inference in the connectome with permutation-based FWER correction (2), 4) **“cluster (TFCE)”**: Threshold-Free NBS (tfNBS), a threshold-free method for cluster-level inference with permutation-based FWER correction (17), 5) **“network”**: the Constrained NBS (cNBS) method we introduced for network-level inference (18) with permutation-based estimation of nulls followed by parametric FWER correction (Bonferroni procedure), 6) **“network (FDR)”**: the cNBS method with FDR correction (Simes procedure), and 7) **“whole brain”**: the Multivariate cNBS statistic (mv-cNBS) we introduce here for whole-brain multivariate inference based on cNBS. All procedures used nonparametric permutation-based procedures for estimating and controlling FWER except the edge-level approaches. The Shen268 atlas was also used to define 10 node communities (cf. (39) for partitioning the graph for cNBS and mv-cNBS. All inferential procedures have been implemented as extensions to the Matlab NBS toolbox (except the NBS procedure, for which we used the original procedure implemented in the toolbox). The tool was also extended to use procedures from the Matlab command line.

All code for inference, benchmarking, and summarization used here is available at: https://github.com/SNeuroble/NBS_benchmarking. See **SI Methods** for details.

## Supporting information

Supplemental Methods and Results

## Author contributions

S.N. conceived of and designed the project, wrote the code, and performed analyses. D.S. contributed to the design of the project and provided supervision. A.F.M. proposed the inclusion of a multivariate approach and provided feedback during the project. A.Z. proposed the inclusion of community randomization tests and provided feedback during the project. S.N. wrote the manuscript with input from D.S., A.F.M., and A.Z.

## Acknowledgments

This work was supported by the National Institute of Mental Health under award number K00MH122372 (S.N.) and by the National Institute of Biomedical Imaging and Bioengineering under award number R01EB027119 (A.F.M). Data were provided by the Human Connectome Project, WU-Minn Consortium (Principal Investigators: David Van Essen and Kamil Ugurbil; 1U54MH091657) funded by the 16 NIH Institutes and Centers that support the NIH Blueprint for Neuroscience Research; and by the McDonnell Center for Systems Neuroscience at Washington University (40). We also thank the Yale Center for Research Computing, specifically Robert Bjornson, for assistance and maintenance of the Farnam cluster.

**Supplemental Materials: Methods, Results, Figures**

## References

1. K. J. Friston, A. Holmes, J. B. Poline, C. J. Price, C. D. Frith, Detecting activations in PET and fMRI: levels of inference and power. Neuroimage 4, 223–235 (1996).

2. A. Zalesky, A. Fornito, E. T. Bullmore, Network-based statistic: identifying differences in brain networks. Neuroimage 53, 1197–1207 (2010).

3. H. R. Cremers, T. D. Wager, T. Yarkoni, The relation between statistical power and inference in fMRI. PloS one 12, e0184923 (2017).

4. S. Noble, D. Scheinost, R. T. Constable, Cluster failure or power failure? Evaluating sensitivity in cluster-level inference. Neuroimage 209, 116468 (2020).

5. S. Gao, G. Mishne, D. Scheinost (2020) Poincaré embedding reveals edge-based functional networks of the brain. in International Conference on Medical Image Computing and Computer-Assisted Intervention (Springer), pp 448–457.

6. J. Faskowitz, F. Z. Esfahlani, Y. Jo, O. Sporns, R. F. Betzel, Edge-centric functional network representations of human cerebral cortex reveal overlapping system-level architecture. Nature neuroscience 23, 1644–1654 (2020).

7. L. Q. Uddin, B. T. Yeo, R. N. Spreng, Towards a universal taxonomy of macro-scale functional human brain networks. Brain topography 32, 926–942 (2019).

8. D.-E. Meskaldji et al., Improved statistical evaluation of group differences in connectomes by screening–filtering strategy with application to study maturation of brain connections between childhood and adolescence. NeuroImage 108, 251–264 (2015).

9. M. J. McKeown et al., Analysis of fMRI data by blind separation into independent spatial components. Human brain mapping 6, 160–188 (1998).

10. Q. H. Lin, J. Liu, Y. R. Zheng, H. Liang, V. D. Calhoun, Semiblind spatial ICA of fMRI using spatial constraints. Human brain mapping 31, 1076–1088 (2010).

11. W. Zhao et al., Individual differences in cognitive performance are better predicted by global rather than localized BOLD activity patterns across the cortex. Cerebral Cortex 31, 1478–1488 (2021).

12. A. Eklund, T. E. Nichols, H. Knutsson, Cluster failure: why fMRI inferences for spatial extent have inflated false-positive rates. Proceedings of the National Academy of Sciences, 201602413 (2016).

13. R. A. Poldrack et al., Scanning the horizon: towards transparent and reproducible neuroimaging research. Nat Rev Neurosci 18, 115–126 (2017).

14. D. Szucs, J. P. Ioannidis, Sample size evolution in neuroimaging research: An evaluation of highly-cited studies (1990–2012) and of latest practices (2017–2018) in high-impact journals. NeuroImage 221, 117–164 (2020).

15. J. M. Bland, D. G. Altman, Multiple significance tests: the Bonferroni method. BMJ 310, 170 (1995).

16. J. D. Storey, A direct approach to false discovery rates. Journal of the Royal Statistical Society: Series B (Statistical Methodology) 64, 479–498 (2002).

17. S. M. Smith, T. E. Nichols, Threshold-free cluster enhancement: addressing problems of smoothing, threshold dependence and localisation in cluster inference. Neuroimage 44, 83–98 (2009).

18. S. Noble, D. Scheinost, The constrained network-based statistic: a new level of inference for neuroimaging. Medical Image Computing and Computer Assisted Intervention (2020).

19. R. J. Simes, An improved Bonferroni procedure for multiple tests of significance. Biometrika 73, 751–754 (1986).

20. Y. Benjamini, Y. J. J. o. t. R. s. s. s. B. Hochberg, Controlling the false discovery rate: a practical and powerful approach to multiple testing. 57, 289–300 (1995).

21. E. B. Wilson, Probable inference, the law of succession, and statistical inference. Journal of the American Statistical Association 22, 209–212 (1927).

22. A. M. Winkler, G. R. Ridgway, M. A. Webster, S. M. Smith, T. E. Nichols, Permutation inference for the general linear model. Neuroimage 92, 381–397 (2014).

23. C.-W. Woo, A. Krishnan, T. D. Wager, Cluster-extent based thresholding in fMRI analyses: pitfalls and recommendations. Neuroimage 91, 412–419 (2014).

24. D. Scheinost et al., Ten simple rules for predictive modeling of individual differences in neuroimaging. Neuroimage 193, 35–45 (2019).

25. J. D. Rosenblatt, L. Finos, W. D. Weeda, A. Solari, J. J. Goeman, All-resolutions inference for brain imaging. Neuroimage 181, 786–796 (2018).

26. J. Chumbley, K. Worsley, G. Flandin, K. Friston, Topological FDR for neuroimaging. Neuroimage 49, 3057–3064 (2010).

27. D. Kessler, M. Angstadt, C. S. Sripada, Reevaluating “cluster failure” in fMRI using nonparametric control of the false discovery rate. Proceedings of the National Academy of Sciences 114, E3372–E3373 (2017).

28. D. van der Meer et al., Understanding the genetic determinants of the brain with MOSTest. Nature communications 11, 1–9 (2020).

29. A. Subramanian et al., Gene set enrichment analysis: a knowledge-based approach for interpreting genome-wide expression profiles. Proceedings of the National Academy of Sciences 102, 15545–15550 (2005).

30. S. M. Smith et al., Correspondence of the brain’s functional architecture during activation and rest. Proceedings of the national academy of sciences 106, 13040–13045 (2009).

31. M. W. Cole, D. S. Bassett, J. D. Power, T. S. Braver, S. E. Petersen, Intrinsic and task-evoked network architectures of the human brain. Neuron 83, 238–251 (2014).

32. X. Di, S. Gohel, E. H. Kim, B. B. Biswal, Task vs. rest—different network configurations between the coactivation and the resting-state brain networks. Frontiers in human neuroscience 7, 493 (2013).

33. J. R. Cohen, M. D’Esposito, The segregation and integration of distinct brain networks and their relationship to cognition. Journal of Neuroscience 36, 12083–12094 (2016).

34. E. M. McCormick, K. L. Arnemann, T. Ito, S. J. Hanson, M. W. Cole, Latent functional connectivity underlying multiple brain states. bioRxiv (2021).

35. A. S. Greene, S. Gao, S. Noble, D. Scheinost, R. T. Constable, How tasks change whole-brain functional organization to reveal brain-phenotype relationships. Cell reports 32, 108066 (2020).

36. S. Marek et al., Towards Reproducible Brain-Wide Association Studies. bioRxiv (2020).

37. M. Salehi, A. Karbasi, D. S. Barron, D. Scheinost, R. T. Constable, State-specific individualized functional networks form a predictive signature of brain state. bioRxiv, 372110 (2018).

38. X. Shen, F. Tokoglu, X. Papademetris, R. T. Constable, Groupwise whole-brain parcellation from resting-state fMRI data for network node identification. Neuroimage 82, 403–415 (2013).

39. S. Noble et al., Influences on the test–retest reliability of functional connectivity MRI and its relationship with behavioral utility. Cerebral Cortex 27, 5415–5429 (2017).

40. D. C. Van Essen et al., The WU-Minn Human Connectome Project: an overview. Neuroimage 80, 62–79 (2013).

