## Supplemental Figures for "Leveling up: improving power in fMRI by moving beyond cluster-level inference"

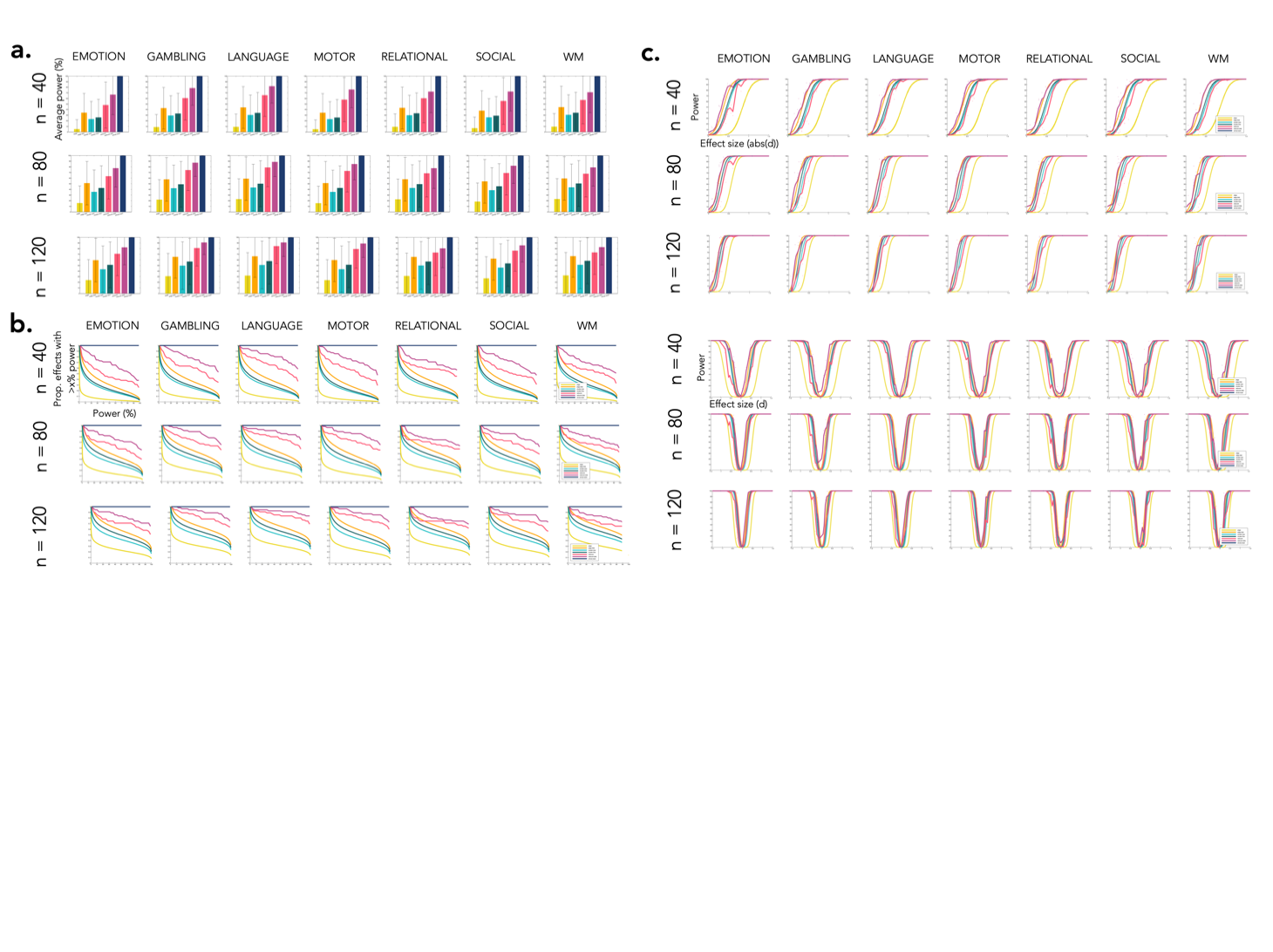

**Supplemental Fig. 1**. **Power across levels of inference by task.** Results are shown for each inferential procedure at three sample sizes and for each of the seven tasks. a) Power to detect an average effect. Error bars represent standard deviation across effects for that task. b) Proportion of effects exceeding each power level. c) Relationship between power and effect size at each level. For all plots, the commonly desired 80% power threshold is indicated by the grey dashed line. Top shows results for absolute value of effect size, bottom for signed effect size.

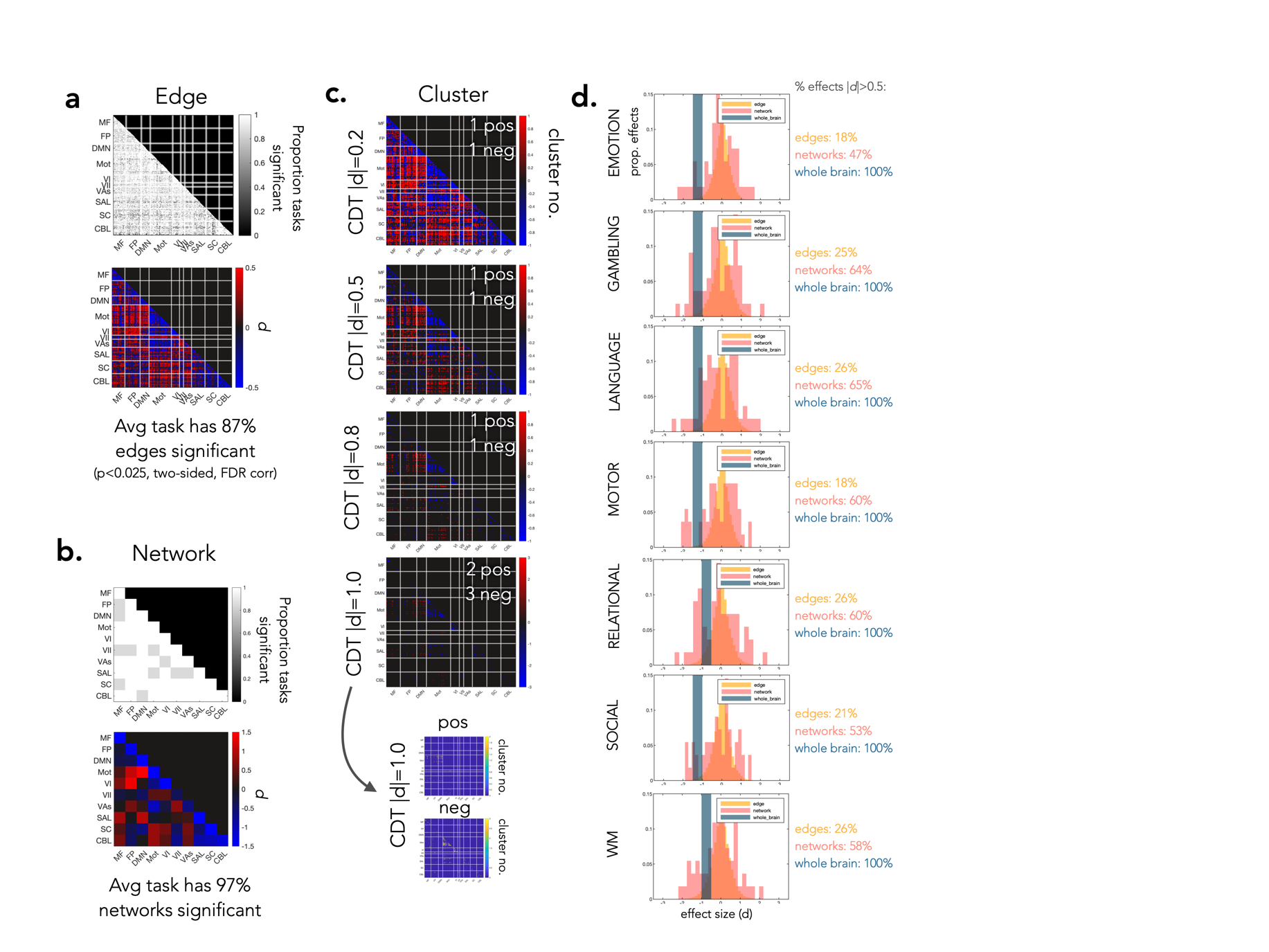

**Supplemental Fig. 2**. **Details of spatial extent of effects in the full “ground truth” dataset.** a) Edge-level effect sizes and proportion of edges marked significant across tasks (p<0.05, two-sided t-test, FDR corrected). b) Network-level effect sizes and proportion of networks marked significant across tasks (p<0.05, two-sided t-test, FDR corrected). Within-community connectivity, generally lower during task than rest, is highlighted by the yellow dotted rectangle. c) Clusters extents across cluster determining thresholds. All edges surviving a cluster-determining threshold of |d|=0.2 (small; i.e., edges with *d*>0.2 and *d*<-0.2), 0.5 (medium), 0.8 (large), and 1.0 (very large) are shown and clusters of contiguous edges are counted. Only a single very large cluster determining threshold (|d|=1.0) yields more than one cluster. The bottom pair of figures shows the two positive and three negative clusters reported at that threshold, which are mainly bound by the predefined network definitions. d) Histograms of effect size at the edge-level (40 bins), pooled within networks (20 bins), and pooled across the whole brain (2 bins).

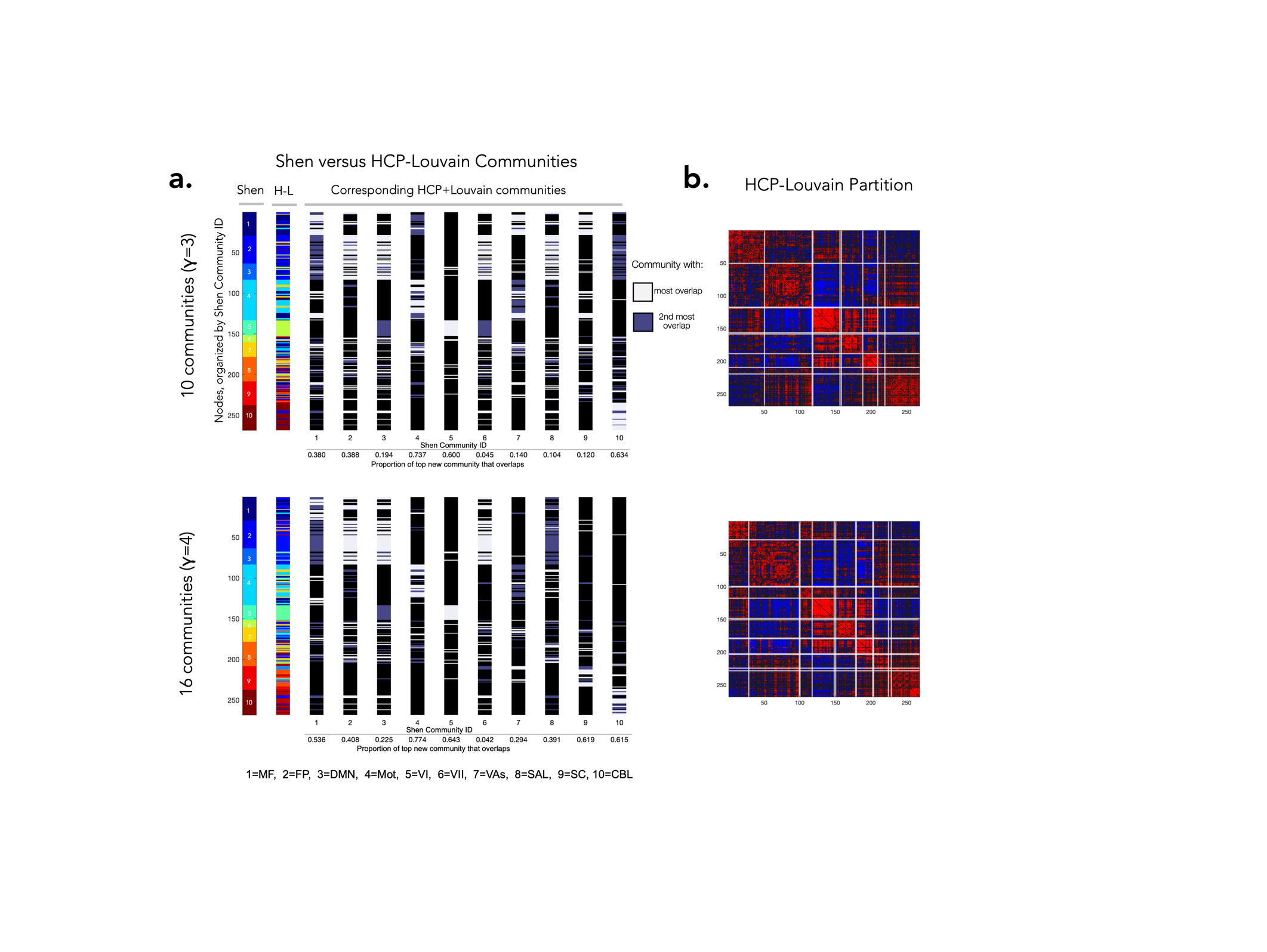

**Supplemental Fig. 3**. **Generalizability of the Shen communities to communities estimated in the HCP data via the Louvain algorithm.** Two parameters were used to estimate communities from the rest-mean task contrast: γ=3 (top; larger communities) and γ=4 (bottom; smaller communities). a) For each bar, nodes are presented in the same order, grouped by the Shen 10 communities. The Shen communities are shown first, followed by the HCP Louvain communities. Subsequent bars to the left show the HCP-Louvain community which best overlaps with each Shen communities, defined as the community with the most nodes within the respective community. For example, the bar for Shen community #1 shows in white the nodes of the HCP-Louvain community that most overlap with Shen community #1. The second most overlapping community is also provided in dark grey. b) HCP-Louvain graph partition based on reordering nodes into the newly estimated communities.

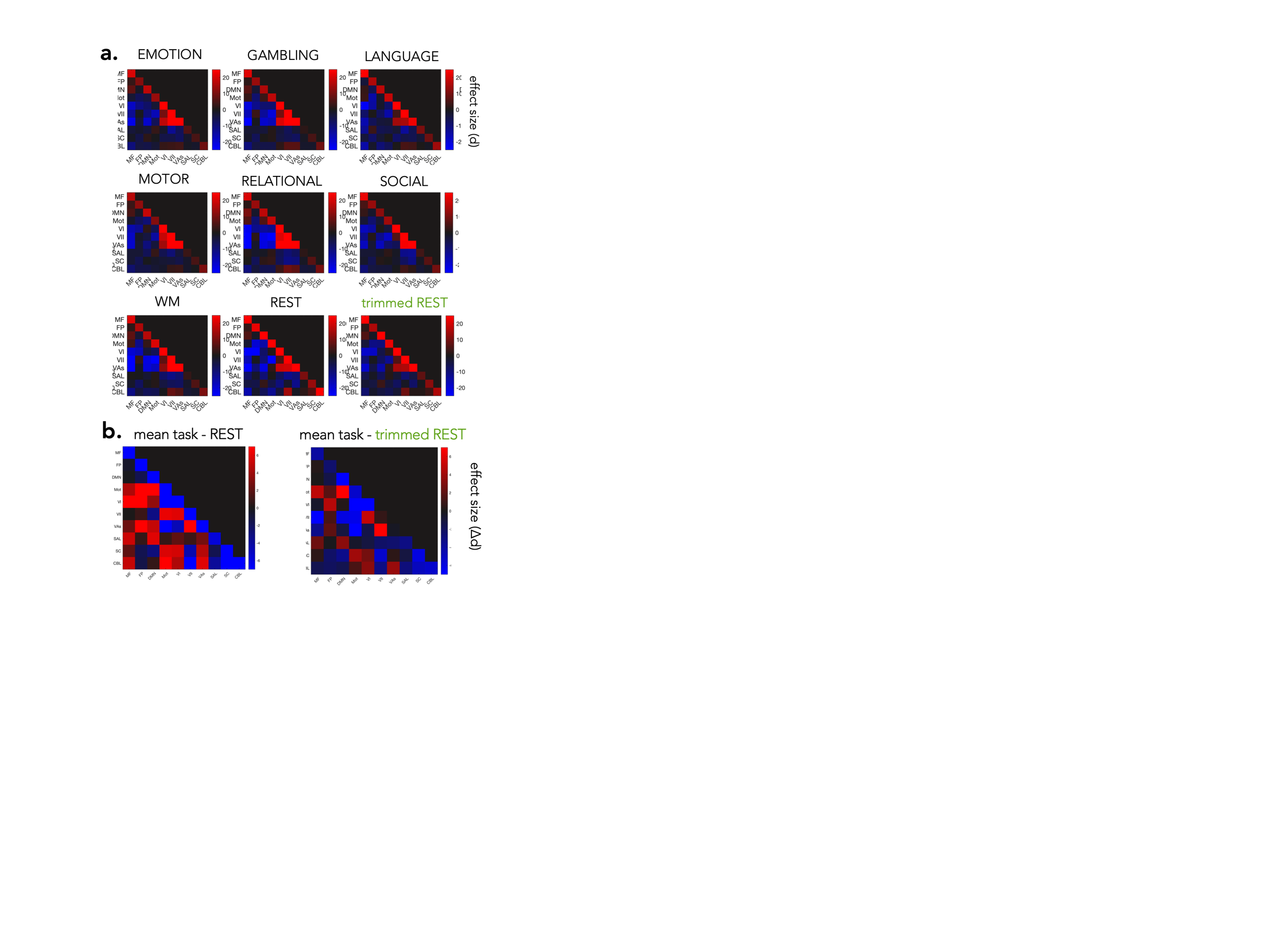

**Supplemental Fig. 4**. **Comparison between all tasks and rest, both original and trimmed.** a) Effect size of each scan. b) Difference between mean effect size across all tasks and rest effect size. Note that Δ*d* for mean(task) *d* - rest *d* shown here are not the same as the mean *d* of the paired contrast between task and rest in the main text. For all plots, effects are calculated first at the edge level then averaged within networks. For trimmed rest results, rest was trimmed to match the shortest task (176 frames).

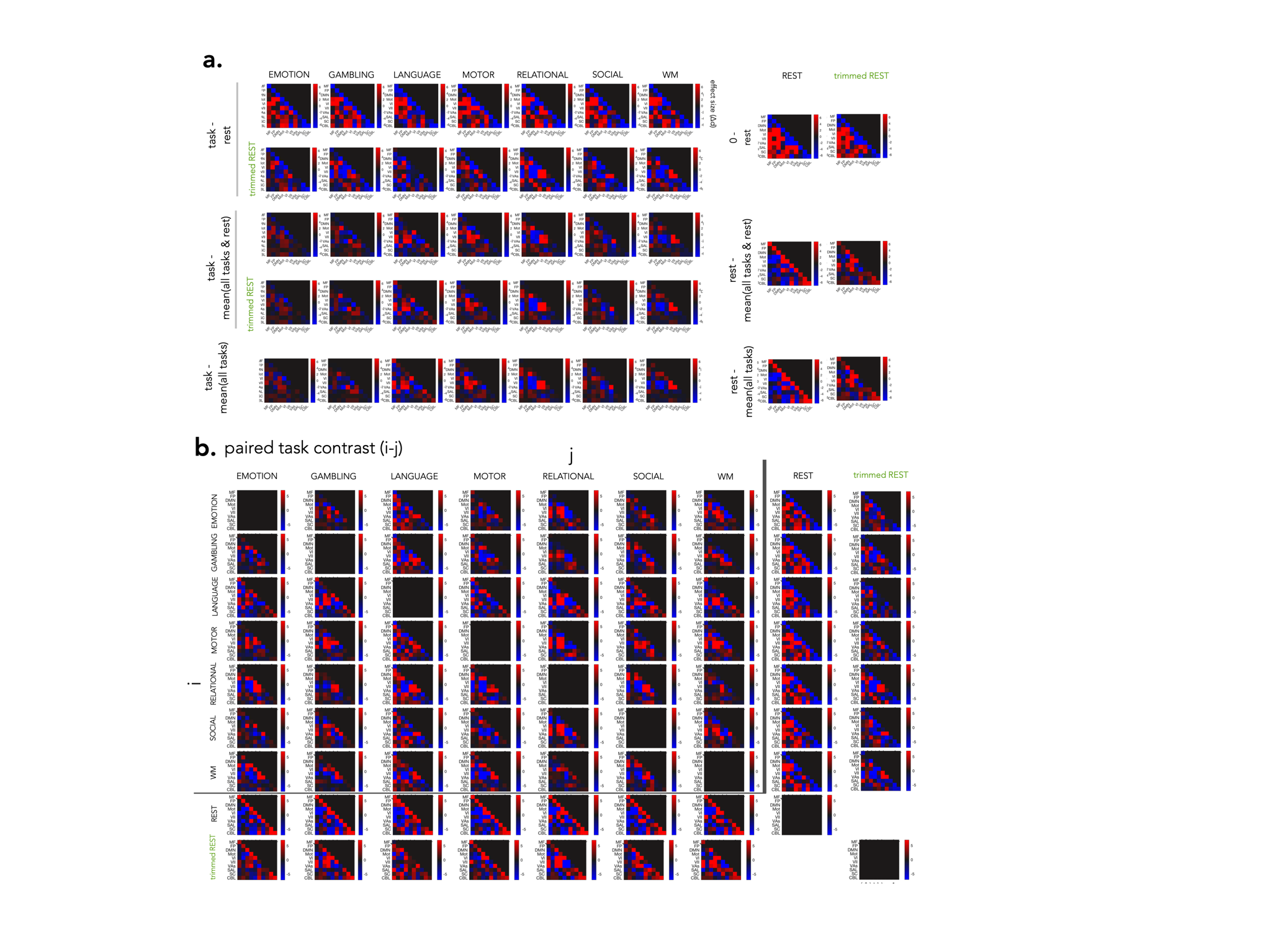

**Supplemental Fig. 5**. **Detailed comparison between all tasks and rest, both original and trimmed.** a) Difference between each scan condition (task and rest) and various reference scans. Top, task minus rest. Middle, task minus mean of all tasks and rest. Bottom, task minus mean of all tasks, not including rest. Rows using trimmed rest noted in green. The right two columns show the difference between rest and various references. The same reference scans are used as above, except for the top row, which shows 0 minus rest. b) Differences between all paired scan conditions. Differences are i - j, with i on the vertical axis and j on the horizontal axis. All effects are calculated first at the edge level then averaged within networks. For trimmed rest, rest was trimmed to match the shortest task (176 frames).

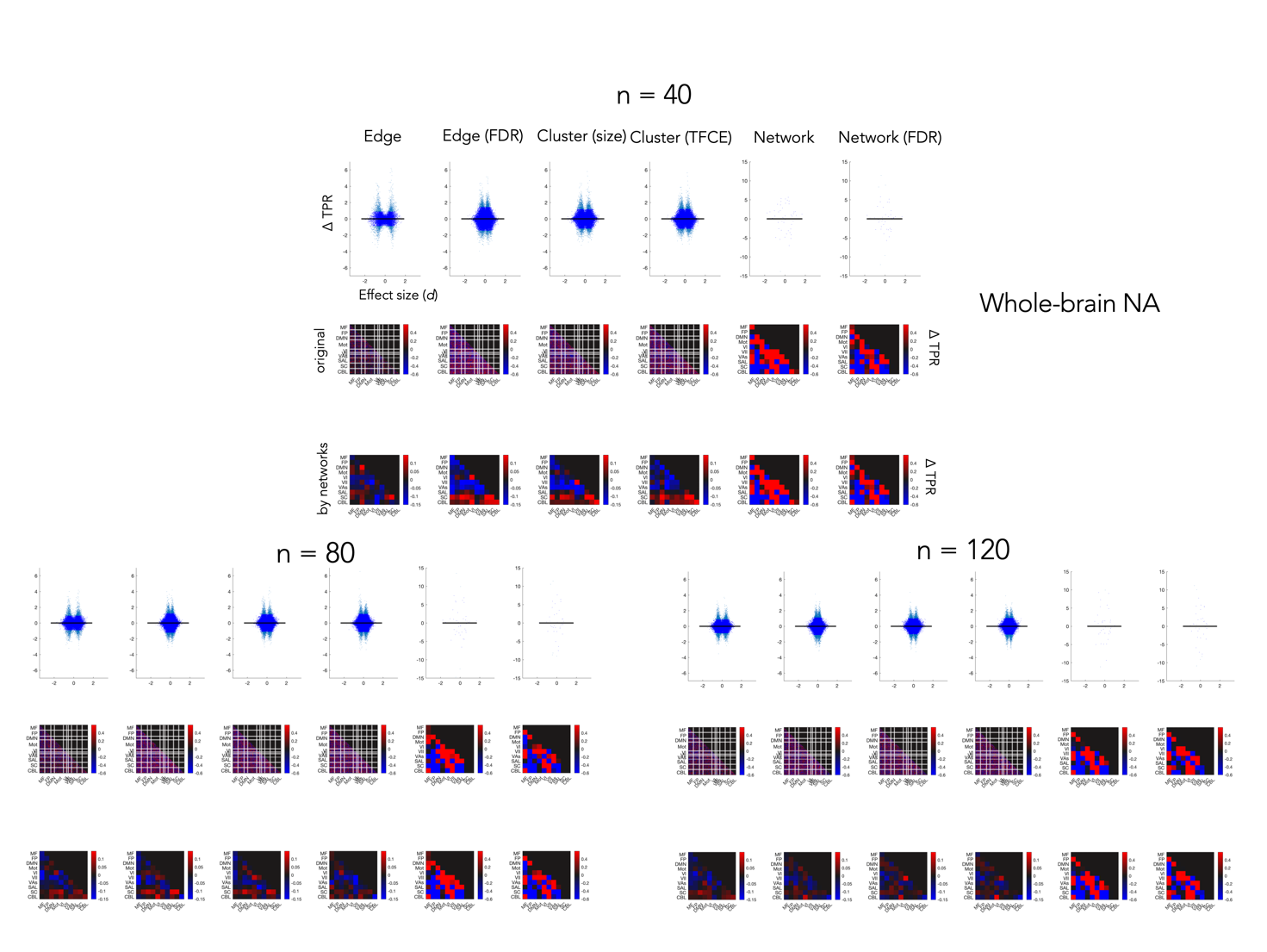

**Supplemental Fig. 6. Spatial bias in power.** Spatial and scatter plots show average task residuals of the effect size versus power curve. For each group size, the top row shows the relationship between residuals and effect size, the middle row shows the effects at the respective level of inference for each inferential procedure, and the bottom row shows the same results averaged within network. The distribution of results is not shown for the full brain procedure since there is only one data point (100% power for each group size).

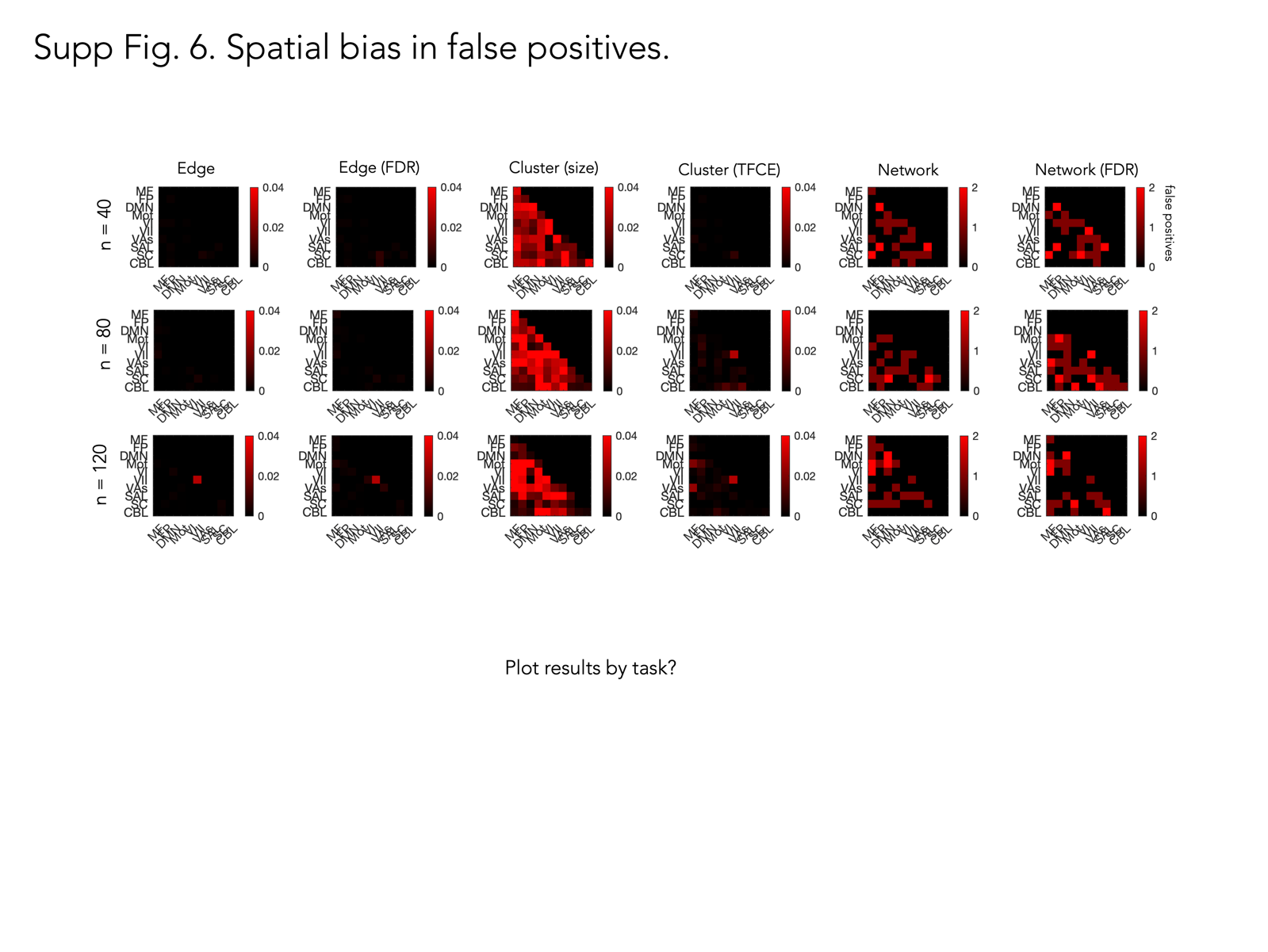

**Supplemental Fig. 7**. **Spatial bias in false positives.** Number of false positives for each inferential procedure, averaged within each network. Rows show results for each group size. The distribution of results is not shown for the full brain procedure since there is only one data point (0 false positives for each group size).
