## Supplemental Methods and Results for "Leveling up: improving power in fMRI by moving beyond cluster-level inference"

The following details the inferential procedures under evaluation (**SI Methods** **1**), data used for benchmarking and preprocessing strategy (**SI Methods** **2**), and benchmarking procedure (**SI Methods** **3**). All code was written in Matlab. To the best of our knowledge, the reporting in this manuscript was consistent with the guidelines provided by the Committee on Best Practice in Data Analysis And Sharing (COBIDAS; see **SI Methods** **4** for details).

**1. Description of Inferential Procedures**

Seven procedures were used to perform inference at each of the four levels, referred to here as “edge”, “edge (FDR)”, “cluster size”, “cluster TFCE, “network”, “network (FDR)”, and “whole brain”. All procedures rely on a user-specified GLM to estimate univariate (edge-level) statistics and specify a target familywise error rate (FWER) or false discovery rate (FDR) of 5%. All procedures are nonparametric except the edge-level procedures and use K=1,000 permutations to estimate the null using the procedure implemented in the NBS toolbox (1). For each permutation, rest and task data are randomly exchanged for each subject, edge-level statistics are calculated, then the null statistic for the given inferential procedure is estimated. This procedure has been shown to offer weak but exact (on average) control of the desired FWER level with a per-repetition 95% confidence interval corresponding with K=1,000 of FWER = 0.0500 ± 0.0138 (<https://fsl.fmrib.ox.ac.uk/fsl/fslwiki/Randomise/Theory>).

All procedures were implemented for the present study as extensions to the NBS toolbox except the “cluster” (NBS) procedure, which used the procedure originally implemented in the toolbox. The toolbox was also modified to be run from the Matlab command line rather than as a toolbox-specific GUI.

**1.1. Edge Procedure**

The “edge” procedure used the Bonferroni algorithm to correct edge-level p-values. The parametric Bonferroni procedure is intended to control the FWER to a specified level and has been previously described in detail by (2).

**1.2. Edge (FDR) Procedure**

The “edge (FDR)” procedure used the Storey algorithm as implemented in the Matlab tool *mafdr* to correct edge-level p-values. The parametric Storey procedure is intended to control the FDR to a specified level and has been previously described in detail by (3). Note that the FDR is expected to control the FWER at the specified level when all tests are drawn from the null (4).

**1.3. Cluster Size Procedure**

The “cluster size” procedure used the Network-Based Statistic (NBS) algorithm implemented in the Matlab NBS toolbox version 1.2 ((1); <https://www.nitrc.org/projects/nbs>). In brief, at each of K permutations, a cluster-determining threshold (here, *t*=3.1, corresponding with *p*=0.005-0.001 for DOF=10-1,000) is applied and each group of contiguous edges (i.e., edges that are connected by a node) is identified as a separate cluster (i.e., “component”). The present study used cluster size as the target statistic, which reflects the number of contiguous edges in a cluster. The maximum cluster size is then recorded at each permutation, and this set of maxima forms the null for inference. That is, significant target (unpermuted) data clusters are determined by comparing the sizes of all target clusters with the cluster sizes in the null.

As an alternative to cluster size, one may also use cluster intensity as the target statistic. Both are implemented in the NBS toolbox. Cluster intensity uses the sum of the edge-level statistics within the cluster as the target statistic. Preliminary testing yielded very similar results, so size was chosen for all NBS analyses presented in this study.

**1.4. Cluster TFCE Procedure**

The “cluster TFCE” procedure implemented here is an analogue of threshold-free cluster enhancement in the task-based activation literature (TFCE; (5)) and echoes other TFCE methods for NBS described in the literature (6, 7). This procedure follows the cluster size procedure described above except the TFCE statistic is used as the target statistic. The TFCE statistic is defined for each edge x as:

$$TFCE\left( x \right)=\int_{{h=h}_{0}}^{h_{x}} {e(h)}^{E}h^{H} dh$$

where *h* is the magnitude (i.e., strength) of the edge-level statistic (*h_0_* is typically zero and *h_x_* is the magnitude at edge *x*), *e* is the cluster extent of all surrounding edges ≥ *h*, and *E* and *H* are constant weighting parameters for *e* and *h.* Put another way, the TFCE statistic for an edge is essentially a weighted sum of surrounding edges in the cluster containing that edge, where clusters are defined by a CDT of 0 and the tops of nearby peaks are discounted from the sum. In contrast, inference based on cluster size relies on the user to set the minimum meaningful effect size, which can lead to results that are arbitrarily threshold-dependent.

As in the activation context (5), the values of *E* and *H* are determined empirically. The present study used the default parameters of *H*=3 and *E*=0.4 determined for the *Mrtrix* software ((8); see <https://github.com/MRtrix3/mrtrix3/blob/master/cmd/connectomestats.cpp>) based on (9), although the developers caution that the optimization of these parameters may not be precise and that results are not robust to small perturbations in the parameters. That said, these parameters are also close to one set of values recommended by in ((6); *H*=[2.25-3] and *E*=0.5; the other recommended set is *H*=[3-3.5] and *E*=0.75), although that study suggests suboptimal power with *E*<0.5 so the present value of *E* may be suboptimal.

While the other levels of inference controlled FDR as the “better” procedure, to our knowledge there is not an available nonparametric procedure for controlling cluster-level FDR. Thus we selected the TFCE procedure as the “better” cluster-level procedure since it has been previously demonstrated to improve power compared with cluster size.

**1.5. Network Procedure**

The “network” procedure used the Constrained Network-Based Statistic (cNBS) algorithm we previously implemented as an extension to the NBS toolbox (10). The algorithm is described in detail in (10); in brief, cNBS aggregates edge-level statistics within predefined networks rather than using clusters defined by the data at hand, estimates the null for each network via permutation, then uses a parametric algorithm to control a specified false positive rate. The “network” procedure in this section used Bonferroni correction to control the FWER. Thus cNBS is a mixed parameteric-nonparametric approach.

The present study used the Shen268 268-node, 10-community partition defined using the Yale High-Resolution Controls dataset (n=40; (11); <https://fcon_1000.projects.nitrc.org/indi/retro/yale_hires.html>). Just as voxels were grouped into 268 nodes using the Normalized Cut procedure as described previously (11), this partition was created by grouping nodes into 10 communities using the Normalized Cut procedure, without requiring nodes within a community to be spatially contiguous. The 10 communities of nodes are used to define 10 x 11 / 2 = 55 unique networks representing connections within and between all communities; these 55 networks are used in cNBS for inference.

**1.6. Network (FDR) Procedure**

The “network (FDR)” procedure used the same cNBS algorithm as above, except the Simes procedure (12) (developed for FDR control by Benjamini-Hochberg; (4)) was used to control FDR. Note that “network FDR” uses a different FDR controlling algorithm than “edge FDR”, which used the Storey algorithm. While the Storey algorithm is expected to be more powerful (3), the Simes algorithm was chosen for network FDR because it is faster as it does not use bootstrapping to estimate the null, and some have suggested it may yield nearly equivalent results as the Storey algorithm in practice (13).

**1.7. Whole Brain Procedure**

The “whole brain” procedure used mv-cNBS, introduced for the first time here as a multivariate analogue of cNBS that integrates information across the whole brain. Mv-cNBS uses the vector of cNBS statistics as the test-statistic. The distribution of null vectors is then estimated via permutation. Finally, deviations of null vectors from the null centroid as measured by Euclidean distance are compared with the deviation of target-statistic vector from the null centroid to get a p-value. Since the full vector is used as a test statistic this procedure does not require multiple testing correction.

We introduce a couple considerations for the use of mv-cNBS. Mv-cNBS can be thought of as a form of omnibus test (i.e., indicating whether there is a departure from the null at at least one of the network-level test statistics). One implicit assumption is that all variables have equal variance and no covariance, such that the same magnitude test statistic excursion in any direction will result in the same p-value and all variables contribute equally to the variance of the test statistic. This is unlikely strictly true; an important example is that smaller networks may be more variable since they average over fewer edges. While we assume this is not a problematic assumption, it may be helpful to estimate covariance (from factors such as network size or the permutation-based null) for whitening.

Second, the null centroid is currently used as the reference for estimating distance. Re-centering at the null is not done in a univariate permutation test (e.g., a reference of zero is used for one-sample tests). While re-centering would not affect p-value estimates for a univariate test since rankings would remain the same, it can affect rankings for this multivariate test (i.e., the smallest data point in the ranking will be that closest to the reference data point, which could change substantially with choice of reference). The most important flaw with this definition is that if all null data points are sufficiently far from zero, then a zero test statistic (e.g., no effect at any network) could be seen as significant. In practice, sufficient permutations should yield a centroid close to zero, although this can decrease with more variables and fewer repetitions. Note that we were able to achieve expected control of FWER, suggesting that the limitations discussed above may not lead to inflated false positives at least when the null is true everywhere.

**2. Data and preprocessing strategy**

An overview of key data characteristics and benchmarking parameters are given in **SI Table 1**.

**2.1. Data description**

The present study used data from the Human Connectome Project S1200 release (14). Data acquisition parameters have been described in detail elsewhere (15); in brief functional data were acquired with a slice-accelerated multiband gradient-echo echo planar imaging (EPI) sequence on a 3T Siemens Skyra (TR=720 ms, TE=33.1 ms, flip angle=52 degrees, resolution=2 mm^3^, multiband factor=8). All available task data in volume space were included. No exclusion criteria were used and data were not stratified by family (see **SI Results: Limited generalizability of “ground truth” estimates of effect size**), although participants who did not have both scan conditions for a given paired sample contrast (e.g., both EMOTION and REST1 data for EMOTION-REST1) could not be selected for that contrast. No task regression was used; this can be thought of as retaining task activation-specific effects (16). The present use of publicly available, de-identified data from the Human Connectome Project and sharing of analysis results has been reviewed and designated as exempt (Exemption 4) by the Yale University Institutional Review Board.

| **Category** | **Description** |
| --- | --- |
| Dataset | Human Connectome Project S1200 release (N = 1200 subjects) |
| Resampling sample size | n = 40; n = 80; n = 120 |
| Task contrasts | **7 tasks for estimating power (task-versus-REST1 contrast):**  EMOTION (N=1022; 176 frames) — shortest task  GAMBLING (N =1057; 253 frames)  LANGUAGE (N =1021; 316 frames)  MOTOR (N =1058; 284 frames)  RELATIONAL (N =1016; 232 frames)  SOCIAL (N =1027; 274 frames)  WORKING MEMORY (WM; N =1058; 405 frames) — longest task  **1 “fake task” for estimating FWER (REST1-versus-REST2, shuffling labels for each subject at each repetition)**  REST1 (N =1067; 1200 frames)  REST2 (N =1011; 1200 frames) |
| Resampling repetitions | R = 500 |
| Test type | Paired-sample, one-sided |
| Inferential procedure | **edge:** Bonferroni procedure for FWER control (parametric)  **edge (fdr):** Storey procedure for FDR control (parametric)  **cluster:** Network-Based Statistic (NBS) procedure with FWER control  **cluster tfce:** Threshold-Free Cluster Enhancement NBS (tfceNBS) procedure with permutation-based FWER control  **network:** Constrained NBS (cNBS) procedure with mixed nonparametric-parametric (Bonferroni) FWER control  **network (fdr):** Constrained NBS (cNBS) procedure with mixed nonparametric-parametric (Storey) FDR control  **whole brain:** Multivariate cNBS (mv-cNBS) procedure (does not require multiple testing correction) |
| Permutations for nonparametric procedures | K = 1000 |

**Supplemental Table 1. Key methodological details.** Data characteristics and benchmarking parameters.

**2.2. Preprocessing and first-level analysis**

Minimally preprocessed data released by the HCP team were used for estimating connectivity. The minimal preprocessing pipeline has been described previously (17, 18); in brief, this included “gradient unwarping, motion correction, fieldmap-based EPI distortion correction, brain-boundary-based registration of EPI to structural T1-weighted scan, non-linear (FNIRT) registration into MNI152 space, and grand-mean intensity normalization,” spatial smoothing with an “unconstrained 3D Gaussian kernel of FWHM=4mm,” computation of activity estimates from the general linear model (including corrections and confound modeling), then temporal filtration and prewhitening.

Data were then further preprocessed in volume space using legacy BioImage Suite (BIS (19); <https://medicine.yale.edu/bioimaging/suite/>; the modern version of the software is BISWeb; <https://bioimagesuiteweb.github.io/webapp/>). Noise covariates were regressed from the data, including linear, quadratic, and cubic drift, mean white matter signal and cerebrospinal fluid signal (defined by automatic segmentation), and a 24-parameter model of motion (six rigid-body motion parameters, six temporal derivatives, and squared terms; (20)). No volume censoring was used. A Gaussian spatial filter was applied (approximate cutoff frequency=0.12Hz). Connectivity matrices were then calculated via Pearson’s correlation between all mean node timecourses in the Shen268 atlas, followed by Fisher transformation to *z*-scores. Finally, left-to-right and right-to-left encoding connectivity matrices, acquired in subsequent runs for each scan condition, were averaged together for each subject. To perform a follow-up analysis about how scan duration influenced task-rest contrasts, connectivity matrices were also estimated from “trimmed” REST1 data, which matched the scan duration of the shortest task (176 frames in the EMOTION task).

**3. Benchmarking true and false positives**

The present study used the benchmarking procedure we previously introduced in (10) (see for algorithm and details); this builds on our previous work (21) and was inspired by (22, 23). As in (10), resampled data were compared with the full sample “ground truth” dataset to determine whether effects were detected in the direction dictated by the ground truth. This is analogous to a one-sided t-test where tests are the conducted in a hypothesized direction that happens to be “correct”. Each test estimated changes in connectivity due to task compared with rest.

**3.1. Estimation of ground truth effect sizes**

To estimate “ground truth” effect sizes, paired sample contrasts were performed for the full sample of each task versus rest (*N*=1016–1058). There are many ways to estimate task-related effects in functional connectivity; here, we use a simple context independent analysis (i.e., original timeseries without using a task regressor), which may be more phenotypically relevant than a context-dependent analysis (i.e., based on a task regressor; (24)). Edge-level effect sizes were calculated directly from the edge-level *z*-scores for each subject, which represent standardized edge strength. Network-level effect sizes were calculated by first averaging *z*-scores within networks for each subject. Similarly, whole brain-level effect sizes were calculated by averaging *z*-scores across the whole connectome for each subject. Paired one-sample *t*-statistics were then obtained for each variable by fitting a GLM with the NBS toolbox. *T*-statistics were then converted to Cohen’s *d* coefficients used to measure effect size here. The conversion for a one-sample *t*-statistic is $d_{s}=\frac{t}{\sqrt{n}}$, where *d_s_* is the Cohen’s *d* coefficient for the sample, *t* is the *t*-test statistic, and *N* is the sample size (from eqn. 2.5.9. of Cohen, 1988, pp. 72; cf. (21)). For reference, effect size is frequently classified as small *d*=0.2, medium *d*=0.5, and large *d*=0.8 (25).

We have previously noted a couple considerations about this measure of “ground truth” (21). As a reminder, the extent to which effects estimated here will generalize to other task-specific contexts or accurately reflect “true” effects of interest is unknown. However, the HCP is one of the largest task-based datasets collected for fMRI and provides one of the best approximations of empirical effects to which we have access. Second, it is important to recognize that a non-zero effect will necessarily be estimated for each variable, but some effects may be smaller in magnitude than the error of the estimator and therefore small effects (and true positive rates calculated based on them) should be interpreted with caution. Additionally, when interpreting both ground truth and benchmarking, one may prefer to limit interpretation of the estimated ground truth effects as known characteristics pertaining only to a fixed empirical distribution (i.e., only the data used here) rather than estimated characteristics of an unknown distribution. Finally, **SI Methods** **3.3** will discuss how ground truth effects were used for true positive rate calculation and summarization, and why these are not always the same as the statistics used for inference.

**3.2. Resampling procedure**

Data were resampled (i.e., repeatedly subsampled) at each of *R*=500 repetitions. For each repetition, *n*=40, 80, or 120 subjects were randomly selected without replacement from the full dataset of *N* subjects for group-level analysis.

Resampling was conducted across three group sizes chosen to span from “small” to “high” sample sizes compared with typical sample sizes for this field (26, 27). The smallest sample size is higher than typical for the field yet still only just adequately powered (power=80%) to detect a medium-sized effect when using a single uncorrected one-sided t-test (27), and the largest sample size is much larger than typical (27) but again only just adequately powered to detect a small effect when using a single test. Resampling was also conducted for eight different scan contrasts (seven real task contrasts and one “fake” task contrast) and each of the seven inferential procedures.

Each inferential procedure was used to perform paired one-sample, one-sided tests resulting in a map of edges that survived correction. All surviving edges were added to a cumulative summary kept across repetitions. This procedure was repeated switching the sign of the tail used for the one-sided test to accumulate surviving edges from the positive and negative tails in two separate cumulative maps of positive tests. Cumulative positive test results from the real task contrasts were used to calculate true positives (**SI Methods** **3.3**), and per-repetition (i.e., not cumulative) positive tests from the “fake” task contrast were used to calculate false positives (**SI Methods** **3.4**).

**3.3. True positive rate calculation**

Note that the following statistical terminology for defining true and false positive counts and rates are derived from notation by (28), which we used previously (21). For consistency with those volumetric works, we continue to use *V* (previously representing voxel-level counts) to represent counts at edges, networks, and the whole brain.

True positive rates were estimated by comparing detected edges and clusters to “ground truth” edge effect signs and comparing detected networks to “ground truth” network effect signs; whole brain results were obtained without reference to the whole brain “ground truth”. For each edge or network *x*, the true positive rate was calculated as follows. The sign of the ground truth effect at *x* was determined. The corresponding positive- or negative-tail cumulative map of positive tests was selected (this is equivalent to selecting the tail of the one-sided test *a priori* based on the ground truth effect sign at *x*). The value of the selected cumulative map at *x*, reflecting the number of times *x* survived thresholding across all resampling repetitions, was interpreted as the number of “true positives” at *x* (*V_x_*_,1|1_). The true positive rate at *x* (TPR*_x_*) was then calculated as the number of true positives at *x* (*V_x_*_,1|1_) divided by the total number of repetitions (*R*), since all repetitions contained a false null (*V_x_*_,⋅|1_=*R*): $\mathrm{TPR}_{x}\mathbf{=}\boldsymbol{V}_{\boldsymbol{x,}\mathbf{1|1}}\mathbf{/}R$ . The same procedure was used for the calculation of true positives for the whole-brain procedure, except only the positive tail results were used and any result was considered a true positive (i.e., no “ground truth” maps were used), since the test statistic uses only the magnitude of the distance from the null centroid and thus is always positive. The true positive rate for the whole-brain procedure is defined as $\mathrm{TPR}\mathbf{=}\boldsymbol{V}_{\mathbf{1|1}}\mathbf{/}R$. As discussed before (21), a couple conceptual points will be reviewed here.

The ground truth data used for true positive rate calculation matched the level of each inferential procedure except cluster-level inference (excluding the whole brain-level, for which ground truth data was not used). This is because a true positive cluster is defined as a cluster where the null was rejected at at least on edge. Thus, for each edge, we can determine how many times it was part of a cluster for which the null was rejected. In contrast, rejecting the null for a cluster does not imply that the null is rejected for any particular edge in that cluster. Rejecting the null for a cluster only implies that the null is rejected for at least one edge in that cluster, and rejecting the null for an edge based on the TFCE statistic implies that the null is rejected for some cluster containing that edge (5). In the activation-mapping context, others have synonymously interpreted a voxel belonging to a detected cluster as a true positive when the null is actually true for an voxel (5, 21, 29).

As discussed above, whole brain-level effect sizes were not used for calculating whole-brain true positives; however, they were used for summarizing whole brain-level true positives (e.g., creating effect size versus power plots). Note that the whole brain-level ground truth effect size (pooled across all connectome edges) does not match the statistic used to perform whole brain-level inference (Euclidean distance of cNBS vector from the null centroid). While one option may have been to use Mahalanobis distance as the multivariate generalization of Cohen’s *d*, Mahalanobis distance cannot be directly interpreted like Cohen’s *d* (i.e., with rules of thumb for small, medium, and large) and thus we opted for a more interpretable pooled effect size.

**3.4. False positive rate calculation**

### A “fake task” paradigm was used to estimate false positives. Namely, when subsampling a group of subjects for benchmarking, the REST1 and REST2 scans were randomly shuffled for each subject. Only positive tail results were used for estimating FWER, since the permutation is expected to produce symmetric test results. Any surviving result at any repetition *r* was labelled a false positive. The idea of using shuffled regressors in resting state data to define a fake task contrast where all positives are false was inspired by (23).

### False positives were defined for an edge or network *x* at *r* as *v_rx_*_,1|0_ and for the whole brain at *r* as *v_r_*_,1|0_. The familywise error rate (FWER) was then calculated as the number of times at least one false positive was found at a repetition divided by the total number of repetitions (*R*) since all repetitions contained a true null (*v_x_*_,⋅|0_=*R*). For edges and networks, $\mathbf{FWER =}\sum_{\boldsymbol{r=1}}^{\boldsymbol{R}} \mathbf{[}\sum_{\boldsymbol{x=1}}^{\boldsymbol{X}} \boldsymbol{v}_{\boldsymbol{x,r}\mathbf{,1|0}}\boldsymbol{\geq1 ] /}\boldsymbol{R}$ , where [...] is the Iverson bracket (1 when true, 0 when false) and *X* is the total number of edges or networks. There is only a single member of the family at the whole brain level, so for that procedure “FWER” was defined as the false positive rate $\mathbf{FWER =}\sum_{\boldsymbol{r=1}}^{\boldsymbol{R}} \boldsymbol{v}_{\boldsymbol{r}\mathbf{,1|0}}\boldsymbol{/ R}$.

### 3.5. Resource availability and technical details

Scripts for performing all inferential procedures, benchmarking, summarization, and visualization in Matlab are available at <https://github.com/SNeuroble/NBS_benchmarking>. Altogether, 7 inferential procedures, 8 contrasts, and 3 resampling group sizes were run for benchmarking power and FWER, resulting in a total of 168 experiments. Benchmarking was run on the Farnam High Performance Computing cluster at Yale via the Simple Linux Utility for Resource Management (SLURM) workload manager (<https://docs.ycrc.yale.edu/clusters-at-yale/clusters/farnam/>). Requested resources per experiment included 13 CPUs and 40GB memory on compute nodes from the general partition (Red Hat Enterprise Linux Server 7.9). Repetitions were parallelized within each experiment using Matlab *parfor*. The following were the average run times across tasks for individual experiments (13 parallel jobs, R=500 repetitions and K=1,000 permutations per job): [0.2, 0.4, 0.6] hours for edge, [0.3, 0.5, 0.7] hours for edge FDR, [3.2, 6.7, 11.4] hours for cluster, [6.9, 8.9, 12.7] hours for cluster TFCE, [3.1, 6.2, 10.1] hours for network, [3.2, 8.0, 10.0] hours for network FDR, and [3.1, NA, 11.2] hours for whole brain (intervals represent run time corresponding with for group size *n*=40, *n*=80, *n*=120; note that *n*=80 for whole brain took longer than a day to run for unexplained reasons so we leave that NA). For comparison, sequential run time for a single experiment is estimated to be as little as 2.6 hours (40 subjects, edge) or as much as 165 hours (120 subjects, cluster TFCE).

In addition to the above, experiments were also run to benchmark nonparametric FDR as implemented in the NBS toolbox. By far, nonparametric FDR took the most resources to run, requiring double to triple the memory and time to run as the other approaches. Since FWER control was not achieved with the above number of permutations, nonparametric FDR experiments were also repeated with 10,000 permutations and FWER was still not achieved.

### 4. COBIDAS Compliance

To the best of our knowledge, the present manuscript is compliant with the mandatory COBIDAS recommendations (i.e., those marked Y in the checklist). The following summarizes which categories have and have not been explicitly described in the present manuscript as well as our rationale.

First, there are a number of data description categories that were not described in the present manuscript because they have been previously extensively documented by the Human Connectome Project team. These categories include: Number of subjects (scanned, excluded in database after acquisition), Inclusion criteria and descriptive statistics, Ethical considerations (for data collection centers), Design specifications, Task specification, Behavioral performance, All Acquisition Reporting, Preprocessing Reporting (for minimal processing pipeline), and Reproducibility (first level).

The following were not described in the present manuscript because they were not relevant to the present study: Statistical Modeling & Inference: Predictive analysis, and Results Reporting: Functional connectivity and Multivariate modelling & predictive analysis (these seem relevant but are not; see descriptions).

The categories that uniquely pertain to the present study and are therefore described in the present manuscript include: Number of subjects (excluded and analyzed), Ethical considerations (for local IRB), Power analysis, Preprocessing reporting (further processing beyond HCP minimal pipeline), Statistical Modeling and Inference: Mass univariate analyses, Functional connectivity, and Multivariate modeling, Results reporting: Mass univariate analysis, Data Sharing, and Reproducibility.

**SUPPLEMENTAL RESULTS**

**Limited accuracy of estimated “ground truth” effects**

Here, we treated the full sample as the population of interest in order to form a more realistic (i.e., empirical) estimate of “ground truth” effects for benchmarking. While this choice is expected to provide a reasonable framework for estimating power, this is of course not an exact measure of true population effects. In particular, given the large number of edges, it is likely that many estimated edge-level effects show particularly high errors in magnitude (i.e., some edges are much larger or much smaller than the truth). Sign errors are expected to be more likely for effects measured to be small in magnitude. This concern is partly mitigated by the fact that the majority of edges and networks in the full sample were found to be significant (Fig. 3), implying that the majority of effect signs are likely to be meaningful and thus useful for benchmarking power.

Additionally, there is prominent family structure in the HCP associated with heritable task-based activation profiles (30). We did not use this in the estimation of effects and instead opted for a simple approach that treats all individuals as independent observations from a population. We anticipate that the unaccounted for dependence between individuals may somewhat bias estimates of ground truth effect sizes. Furthermore, the fact that this dependence structure is not matched during resampling may also somewhat bias estimates of power. However, we do not anticipate that accounting for family structure will substantially alter the main findings.

**Generalizability of the Shen268 partition to the HCP data**

The Shen268 10 community atlas defined in the Yale High-Resolution Controls dataset was used to define a partition specifying 55 unique networks (see details in **SI Methods: 1.5**). We estimated whether this definition of networks showed substantially more between-network variability in the present data (HCP) than randomly defined networks. 10,000 permutations were used to test whether there was significant variability amongst 1) all 55 networks compared with randomly reassigning nodes across the 10 communities in the same partition (nodes rather than edges are shuffled in order to preserve the graph structure), 2) the 10 within-community networks compared with randomly shuffling edges across all within-community networks in the same partition, and 3) the 45 between-community networks compared with randomly shuffling edges across all between-community networks in the same partition. Greater variability was observed in all three cases for the original networks (p<1E-10; **Fig. 3d**). These findings are consistent with the idea that the original partition captures some of the structure in the present dataset, and not just because it separates within- and between-community networks.

We also estimated generalizability of the Shen268 partition to a partition derived from the HCP data using the Louvain method for community detection (community_louvain function in the Brain Connectivity Toolbox; (31); <https://sites.google.com/site/bctnet/measures/list#TOC-Clustering-and-Community-Structure>), which essentially forms communities by maximizing the sum of within-community weights. However, since task-rest contrasts generally showed negative edge weights within the Shen268 communities and positive weights between those communities, rest-task contrasts were instead used to facilitate comparison. Note that these edge weights represent *increased* connectivity during rest, rather than more direct measure of association typically used for community detection. Two sets of communities were estimated using two parameters specifying smaller (γ=3) and larger (γ=4) communities, and the “negative_asym” flag was provided to use separate scalings for defining the modularity matrix using positive and negative edge weights. To obtain a stable definition of each set of communities, 10,000 iterations of this procedure were run and the most common assignment for each node was used as the final HCP-Louvain community assignment.

We then compared the overlap between the two sets of communities by 1) calculating which HCP-Louvain community that had the most nodes within each Shen268 community, then 2) calculating the proportion of nodes in that HCP-Louvain community that overlapped with the Shen268 community (**SI Fig. 3**). For the most part there appeared to be fair overlap between the sets of communities. A couple HCP-Louvain communities showed high overlap with a Shen268 community (Motor, V1, cerebellum). There were a few cases that suggested merging multiple of the Shen268 communities (e.g., merging frontoparietal, default mode, and VII), and several cases where there was relatively little unique overlap (e.g., VII and VAs, both some of the smallest communities). Altogether, despite being defined in a different dataset using rest-only data, the Shen268 partition seems to capture some of the structure of the task-related effects in the HCP and network-level pooling may be a meaningful place to start.

**Differences between task and rest, and effect of unbalanced scan durations**

All task and rest scans showed a similar spatial pattern of connectivity (**Fig. 3; SI Fig. 4a**). Connectivity was generally strong and positive within communities and often weaker or negative between communities. Effects associated with visual communities were particularly strong, with more positive connectivity amongst the visual communities and more negative connectivity between visual and other cortical communities. In contrast, subcortical and cerebellar connectivity was generally relatively weak.

The full resting scan showed stronger connectivity overall than any of the tasks, especially within-community (**SI Fig. 4**). However, rest was the longest of the scans (1200 frames), about triple that of the shortest task (176 frames in the EMOTION task). Since scan duration is directly related to the power to estimate connectivity (see also (32)), we estimated how effect sizes would change with rest trimmed to a duration matching the shortest task (176 frames; **SI Fig. 4**; detailed contrasts in **SI Fig. 5**). Resting state scans at either duration generally showed more positive connectivity within community and between motor-visual communities than task. Yet the trimmed rest scan showed weaker connectivity than untrimmed rest, and the pattern of increased connectivity between communities during rest compared with task became less clear with trimmed rest. While we don’t expect this to substantially alter the main findings and rest remains consistently distinct from task even at this shortest of scan durations (compare task-task with task-trimmed rest contrasts in **SI Fig. 5b**), care should be taken to account for differences in the scan duration that can bias contrasts towards the longer scan.

**Nonparametric edge-level FDR correction implemented in the NBS toolbox**

As indicated in the main text, all approaches were expected to control FWER in the weak sense, defined as attaining FWER levels below the upper bound of expected FWER when the null is true everywhere (95% confidence interval for FWER=3-7% for 500 repetitions; (33, 34)). This includes FDR controlling procedures (4). Unfortunately, the nonparametric edge-level FDR controlling procedure implemented in the NBS toolbox is expected to require many more permutations than is feasible for the present study to achieve valid control—namely, K=10^6–10^7 permutations (<https://www.nitrc.org/forum/message.php?msg_id=31971>; <https://www.nitrc.org/forum/forum.php?thread_id=4543&forum_id=3444>. This procedure estimates the null for each edge via permutation, then uses the Simes algorithm (12) for FDR correction. The time is takes to perform a single repetition of the present benchmarking experiment increases substantially with the number of permutations, with K=10^3 requiring 2 hours per repetition and a K=5x10^4 requiring 2 days per repetition. FWER decreased accordingly as expected—from FWER=100% (K=10^3) to FWER=20% (K=5x10^4)—so we expect that valid FWER can be achieved with K=10^6–10^7 permutations. However, this is not feasible in the current setting where each inferential procedure is evaluated with 24 experiments (8 tasks and 3 group sizes) and 500 repetitions per experiment. As a historical note, this inferential procedure was originally designed for use with relatively small networks which required fewer permutations and fewer resources per permutation.

One option for users to avoid invalid FWER control is to set the minimum estimable p-value to 1/K rather than 0 (i.e., by changing line 89 on NBSfdr.m from "pvals=zeros(1,J)" to "pvals=ones(1,J)"). However, this impacts power and it is instead recommended to use sufficient permutations when using the above procedure.

**SUPPLEMENTAL REFERENCES**

1.	A. Zalesky, A. Fornito, E. T. Bullmore, Network-based statistic: identifying differences in brain networks. *Neuroimage* **53**, 1197-1207 (2010).

2.	J. M. Bland, D. G. Altman, Multiple significance tests: the Bonferroni method. *BMJ* **310**, 170 (1995).

3.	J. D. Storey, A direct approach to false discovery rates. *Journal of the Royal Statistical Society: Series B (Statistical Methodology)* **64**, 479-498 (2002).

4.	Y. Benjamini, Y. J. J. o. t. R. s. s. s. B. Hochberg, Controlling the false discovery rate: a practical and powerful approach to multiple testing. **57**, 289-300 (1995).

5.	S. M. Smith, T. E. Nichols, Threshold-free cluster enhancement: addressing problems of smoothing, threshold dependence and localisation in cluster inference. *Neuroimage* **44**, 83-98 (2009).

6.	H. C. Baggio *et al.*, Statistical inference in brain graphs using threshold‐free network‐based statistics. **39**, 2289-2302 (2018).

7.	T. Spisak *et al.*, Probabilistic TFCE: A generalized combination of cluster size and voxel intensity to increase statistical power. *Neuroimage* **185**, 12-26 (2019).

8.	J.-D. Tournier *et al.*, MRtrix3: A fast, flexible and open software framework for medical image processing and visualisation. *Neuroimage* **202**, 116137 (2019).

9.	L. Vinokur, A. Zalesky, D. Raffelt, R. Smith, A. Connelly, A Novel Threshold-Free Network-Based Statistics Method: Demonstration using Simulated Pathology. *Organization for Human Brain Mapping*, 4144 (2015).

10.	S. Noble, D. Scheinost, The constrained network-based statistic: a new level of inference for neuroimaging. *Medical Image Computing and Computer Assisted Intervention* (2020).

11.	X. Shen, F. Tokoglu, X. Papademetris, R. T. Constable, Groupwise whole-brain parcellation from resting-state fMRI data for network node identification. *Neuroimage* **82**, 403-415 (2013).

12.	R. J. Simes, An improved Bonferroni procedure for multiple tests of significance. *Biometrika* **73**, 751-754 (1986).

13.	N. Pike, Using false discovery rates for multiple comparisons in ecology and evolution. *Methods in ecology and Evolution* **2**, 278-282 (2011).

14.	D. C. Van Essen *et al.*, The WU-Minn Human Connectome Project: an overview. *Neuroimage* **80**, 62-79 (2013).

15.	S. M. Smith *et al.*, Resting-state fMRI in the human connectome project. *Neuroimage* **80**, 144-168 (2013).

16.	A. S. Greene, S. Gao, S. Noble, D. Scheinost, R. T. Constable, How tasks change whole-brain functional organization to reveal brain-phenotype relationships. *Cell reports* **32**, 108066 (2020).

17.	M. F. Glasser *et al.*, The minimal preprocessing pipelines for the Human Connectome Project. *Neuroimage* **80**, 105-124 (2013).

18.	D. M. Barch *et al.*, Function in the human connectome: task-fMRI and individual differences in behavior. *Neuroimage* **80**, 169-189 (2013).

19.	A. Joshi *et al.*, Unified framework for development, deployment and robust testing of neuroimaging algorithms. *Neuroinformatics* **9**, 69-84 (2011).

20.	T. D. Satterthwaite *et al.*, An improved framework for confound regression and filtering for control of motion artifact in the preprocessing of resting-state functional connectivity data. *Neuroimage* **64**, 240-256 (2013).

21.	S. Noble, D. Scheinost, R. T. Constable, Cluster failure or power failure? Evaluating sensitivity in cluster-level inference. *Neuroimage* **209**, 116468 (2020).

22.	H. R. Cremers, T. D. Wager, T. Yarkoni, The relation between statistical power and inference in fMRI. *PloS one* **12**, e0184923 (2017).

23.	A. Eklund, T. E. Nichols, H. Knutsson, Cluster failure: why fMRI inferences for spatial extent have inflated false-positive rates. *Proceedings of the National Academy of Sciences*, 201602413 (2016).

24.	A. S. Greene, S. Gao, S. Noble, D. Scheinost, R. T. Constable (2019) How Tasks Change Whole-Brain Functional Organization to Reveal Brain-Phenotype Relationships. in *NEURON-D-19-01606*.

25.	J. Cohen, *Statistical power analysis for the behavioral sciences* (Academic press, 2013).

26.	R. A. Poldrack *et al.*, Scanning the horizon: towards transparent and reproducible neuroimaging research. *Nat Rev Neurosci* **18**, 115-126 (2017).

27.	D. Szucs, J. P. Ioannidis, Sample size evolution in neuroimaging research: An evaluation of highly-cited studies (1990–2012) and of latest practices (2017–2018) in high-impact journals. *NeuroImage* **221**, 117164 (2020).

28.	T. Nichols, S. Hayasaka, Controlling the familywise error rate in functional neuroimaging: a comparative review. *Statistical methods in medical research* **12**, 419-446 (2003).

29.	C.-W. Woo, A. Krishnan, T. D. Wager, Cluster-extent based thresholding in fMRI analyses: pitfalls and recommendations. *Neuroimage* **91**, 412-419 (2014).

30.	Y. Benhajali *et al.*, Subtypes of brain activation are heritable and genetically linked with behavior in the Human Connectome Project sample. (2020).

31.	M. Rubinov, O. Sporns, Complex network measures of brain connectivity: uses and interpretations. *Neuroimage* **52**, 1059-1069 (2010).

32.	J. W. Cho, A. Korchmaros, J. T. Vogelstein, M. Milham, T. Xu, Impact of Concatenating fMRI Data on Reliability for Functional Connectomics. *BioRxiv* (2020).

33.	E. B. Wilson, Probable inference, the law of succession, and statistical inference. *Journal of the American Statistical Association* **22**, 209-212 (1927).

34.	A. M. Winkler, G. R. Ridgway, M. A. Webster, S. M. Smith, T. E. Nichols, Permutation inference for the general linear model. *Neuroimage* **92**, 381-397 (2014).
